# Alterations in chromosome spatial compartmentalization classify prostate cancer progression

**DOI:** 10.1101/2021.04.15.440056

**Authors:** Rebeca San Martin, Priyojit Das, Renata Dos Reis Marques, Yang Xu, Rachel Patton McCord

## Abstract

Prostate cancer aggressiveness and metastatic potential are influenced by gene expression, genomic aberrations, and cellular morphology. These processes are in turn dependent in part on the 3D structure of chromosomes, packaged inside the nucleus. Using chromosome conformation capture (Hi-C), we conducted a systematic genome architecture comparison on a cohort of cell lines that model prostate cancer progression, ranging from normal epithelium to bone metastasis. Here, we describe how chromatin compartmentalization identity (A-open vs. B-closed) changes with progression: specifically, we find that 48 gene clusters switch from the B to the A compartment, including androgen receptor, WNT5A, and CDK14. These switches could prelude transcription activation and are accompanied by changes in the structure, size, and boundaries of the topologically associating domains (TADs). Further, compartmentalization changes in chromosome 21 are exacerbated with progression and may explain, in part, the genesis of the TMPRSS2-ERG translocation: one of the main drivers of prostate cancer. These results suggest that discrete, 3D genome structure changes play a deleterious role in prostate cancer progression.

**Summary:** Through a systematic analysis of chromosome conformation capture in a cohort of cells that model cancer progression, San Martin et.al. find that rearrangement of the 3D genome structure in prostate cancer is a potential mechanism for disease exacerbation and that genome-wide compartment identity can classify cancer according to progression.

## Introduction

Prostate cancer (PCa) is the predominant new cancer diagnosis in males in the United States. It is also the second most common cause of male cancer-related deaths, second only to lung cancer (Siegel et al., 2020). Patients with late-stage prostate cancer present with a higher incidence of metastases to trabecular bone (Jacobs, 1983, Bubendorf et al., 2000, Hernandez et al., 2018). The specific mechanisms that promote metastasis to bone are not understood. However, disseminated tumor cells can be detected in the blood of about 25% of PCa patients with localized disease. The abundance of these circulating tumor cells positively correlates with metastatic occurrence (Moreno et al., 2005, Danila et al., 2007, Todenhofer et al., 2016).

Metastatic cancer cells migrate out of the primary site squeezing through gaps much smaller than their nuclei, such as interstitial spaces within the organ and tight endothelial junctions to gain access to the circulation (Barbazan et al., 2017, Bergeman et al., 2016, Mierke, 2008). The rigidity of the nucleus, which depends on the inherent stiffness of the nuclear lamina and chromosomal ultrastructure, is the limiting step of this migration (Davidson et al., 2014, Hatch and Hetzer, 2016, Lammerding et al., 2006). The human genome’s packaging into the nucleus’s constrained space requires a systematic organization: from chromosomal territories (Meaburn and Misteli, 2007) to transcriptionally active and inactive chromatin compartments (Lieberman-Aiden et al., 2009). Within compartments, chromatin further organizes into topologically associating domains (TADs), which are segregated from each other by the insulator protein CTCF (Dixon et al., 2012, Nora et al., 2017), and finally into chromatin loops (Nuebler et al., 2018). With the advent of technologies such as genome-wide chromosome conformation capture (Hi-C), it has become clear that these structures are essential for proper gene regulation, DNA replication, and repair (Hnisz et al., 2016, Dekker and Mirny, 2016, Pope et al., 2014, McCord and Balajee, 2018). Further, rearrangement of these domains can impact both the nuclei’s ability to squeeze through tight spaces during metastatic migration and the expression patterns of oncogenes (Gerlitz and Bustin, 2010, Stephens et al., 2018, Hnisz et al., 2016, Barutcu et al., 2015). Nuclear atypia is a common diagnostic tool in PCa (Verdone et al., 2015, Diamond et al., 1982), and there is evidence that that the nuclear lamina content of prostate cells differs between normal epithelium, BPH, and cancer (Partin et al., 1993). This suggests that genome architectural changes may occur in prostate cancer and might influence cancer-promoting gene expression profiles and the nuclear malleability necessary for cancer cells to metastasize.

While several genomic loci have been associated with a higher risk for prostate cancer (Ahmadiyeh et al., 2010, Helfand et al., 2015, Du et al., 2016), one of the most predominant features of poor patient prognosis is the TMPRSS2-ERG translocation in chromosome 21 (Zhou et al., 2020, Demichelis et al., 2007, Tomlins et al., 2005). Interestingly, it has been shown that overexpression of ERG results in chromatin conformation changes (Rickman et al., 2012), but whether the gene fusion occurs due to increased transcription remains unclear. Since it is known that relative chromosome proximity can influence the pattern of translocations that occur(Zhang et al., 2012, Balajee et al., 2018), this type of local rearrangement could potentially result alterations in chromosome compartmentalization that increase contact frequency among the different loci, as previously described in other systems (Engreitz et al., 2012).

Recent studies into the 3D genome structure associated with prostate cancer (Rhie et al., 2019, Luo et al., 2017a, Taberlay et al., 2016), have contributed some insight into regulatory chromatin loops, epigenetic alterations, and the influence that variable structures have in transcription. However, these studies did not address whether there are early changes in genome architecture that persist throughout progression. In this context, changes in compartment identity from transcriptionally repressed heterochromatin to euchromatin poised for transcription could identify genes required for early oncogenesis and those necessary for metastasis. Further, changes in TAD positioning, or shifting of TAD boundaries could reveal altered interactions of neighboring promoter-enhancer regions.

In the present study, we use a combination of Hi-C and ChIP-seq to characterize the genome organization across a cohort of nine cell lines that model prostate cancer progression from the normal epithelium to bone metastasis, including two bone metastatic cell lines of African American lineage. We further assess the different hierarchical levels of genome organization: from large inter-chromosomal translocations to compartment identity to TAD location and TAD boundary shifting. We have identified a cohort of 387 genes that change compartment identity across prostate cancer progression. Interestingly, most of these genes switch compartments as proximal clusters. These compartment identity changes are accompanied by distinct structural features at higher resolution such as gain or loss of TAD structure, stalled transcriptional loops and structural deserts, and TAD boundary appearance, disappearance, or positional shifting. Finally, our results revealed several “genomic architecture hotspots” whose structural changes are persistent throughout the metastatic models; these include WNT5A, CDK14, androgen receptor (AR), and the TMPRSS2-ERG locus, among others. These results suggest that the 3D genome structure can be used as a prognostic marker for the progression of prostate cancer to bone metastasis.

## Methods

### Cell lines

RWPE-1, LNCaP, DU145, 22RV1, VCaP and PC3 cell lines were obtained from the Physical Sciences Oncology Network Bioresource Core Facility, supported by ATCC. All cell lines were cultured according to standard protocols, subculturing cells as they reached 80% confluency with the following media formulations: RWPE media was comprised of keratinocyte specific media supplemented with EGF and bovine pituitary extract (Gibco 17005042). RPMI (Gibco 11835030) was supplemented to match the suggested formula by ATCC (4.5 g/L glucose, 2.383 g/L HEPES and 0.11 g/L sodium pyruvate) and 10% FBS (Corning 35-010-CV). This media was used for LNCaP, 22RV1 and PC3. DU145 cells were cultured in DMEM supplemented with 10% FBS, and DMEM F12: Ham 1:1 (Gibco 11-320-033) with 10% FBS was used for VCaP.

Cell lines MDAPCa2a and MDAPCa2b were a kind gift of Dr. Nora Navone (MD Anderson Cancer Center, Houston TX). These cell lines were grown in HPC1 media (Athena Enzyme Systems) supplemented with 10% FBS, on standard cell culture T75 flasks coated with FCN Coating Mix (Athena Enzyme Systems) as per the manufacturer’s instructions.

All media formulations were supplemented with 100 µg/ml penicillin-streptomycin (Gibco 15-140-122).

### Chromosome Conformation Capture (Hi-C)

Cell pellets for chromosome conformation capture were prepared as previously described (Golloshi et al., 2018). Briefly, cells growing in monolayer in standard T75 flasks were quickly washed with 10 ml of HBSS (Gibco 14-025-134) at room temperature and crosslinked with 10 ml 1% formaldehyde (Fisher Bioreagents BP531-25, in HBSS) for 10 min on a shaking platform. The crosslinking reaction was quenched by adding glycine to a final concentration of 0.14 M (MP biomedicals ICN19482591), followed by a 5 min incubation at room temperature, with shaking. After cooling down the plates on ice for 15 min, the formaldehyde solution was aspirated from the plate and substituted with 10 ml of ice cold HBSS, containing 1X Halt Protease Inhibitor cocktail (Thermo PI78438). Five million cell aliquots were collected by centrifugation and snap frozen in liquid nitrogen.

VCaP, MDAPCa2a and PC3 Hi-C was conducted as described (Golloshi et al., 2018), using DpnII (New England Biolabs). Hi-C was also performed on LNCaP, DU145, VCaP, MDAPCa2a, MDAPCa2b and PC3, using the Arima Hi-C (Arima Genomics) kit, following the manufacturer’s protocol A160141 v01 for library amplification using the NEBNext Ultra II kit (NEB E7645S).

Sequencing was performed by Genewiz (South Plainfield, NJ) on either an Illumina NovaSeq or HiSeq platform with 50 or 150 bp paired end reads. Sequenced reads were mapped to a reference human genome (hg19), binned, and iteratively corrected according to established pipelines (Imakaev et al., 2012) using the dekkerlab-cMapping tool available at https://github.com/dekkerlab/cMapping.

In addition, the same analysis above was performed on fastq files from previously published Hi-C datasets for RWPE, 22RV1 and LNCaP C4-2B: GSE118629 and GSE73782 (Rhie et al., 2019, Luo et al., 2017b). See Supplementary Table 1 for all data sources and statistics. Newly generated LNCaP results were checked for consistency with previously published LNCaP Hi-C results (ENCSR346DCU, Taberlay et al., 2016). Since our LNCaP data had a dramatically higher cis/trans ratio than these previously published datasets, comparisons were difficult, and so we proceeded with only our newly generated data.

### Analysis of Hi-C data

All Hi-C data analysis was carried out using the existing cworld-dekker pipeline, available on github (https://github.com/dekkerlab/cworld-dekker) as follows:

Hi-C heatmaps were generated for genome-wide datasets at a 2.5 Mb resolution, and per chromosome at a 250kb resolution, using the heatmap script.

Compartment analysis was performed via principal component analysis on 250 kb binned matrices using the matrix2compartment script. Positive and negative PC1 values were assigned to A and B compartments, respectively.

The PC1 values per bin for the normal epithelial cell line (RWPE) were subtracted from the values from each cancer cell line, resulting in a normalized distribution (referred to as ΔPC1analysis). Significant changes in compartment identity were defined as those bins whose subtracted value fell either under the mean minus 1.5 the standard deviation or the mean plus 1.5 the standard deviation, for at least six cell lines or all four cells lines in the primary axis, as defined by nearest neighbor analysis as described below (SPRING plot).

The genes contained in regions of interest determined from the ΔPC1 analysis were annotated using the knownGene primary table as referenced in UCSC Genome Browser. (https://genome.ucsc.edu/).

To facilitate the visualization of chromatin compartmentalization, heatmaps for cis interactions were generated by first calculating the Z score of the interactions at 250 kb resolution compared to an expected interaction at each distance from the diagonal and then taking the Pearson correlation of each row and column of the heatmap (zScore correlation matrices). Multi compartment track figures and overlays were constructed using the visualization tool Sushi (Phanstiel et al., 2014)

Genome wide topologically associating domain (TAD) boundaries were determined using the matrix2insulation script on 40kb binned matrices, following the insulation score approach with an insulation square size of 500 kb (Crane et al., 2015).

### Calculation of A-A and B-B compartment interaction strengths

To calculate A-A and B-B compartment interaction strengths for each chromosome distance corrected Hi-C intra-chromosomal interaction frequencies at 250 Kb resolution was reordered according to their corresponding PC1 values (from strongest B to strongest A). Then, the reordered intra-chromosomal interaction matrix was smoothed at 500 Kb resolution. Interactions were classified as A-A, B-B and A-B and thresholded to include only the top 20% of interactions. The median value of each A-A, B-B and A-B interactions is calculated. Finally, the relative A-A and B-B compartment interaction strengths were obtained by subtracting the absolute A-B compartment interaction strength from the absolute A-A and B-B interaction compartment strengths respectively.

For the average compartment interaction strength, the mean of the relative A-A and B-B interaction compartment strengths was calculated. A stronger A-A or B-B compartmentalization level compared to the A-B compartment intermixing would produce a higher positive value. On the other hand, a value close to zero suggests a weaker level of compartmentalization.

### SPRING Plot

To construct the SPRING plot of the prostate cancer cells based on the Hi-C compartmental data, the compartment profile of the cells at the 250 Kb resolution was binarized. For example, the genomic regions where the compartment strength are greater than ‘0’, were converted to ‘1’ and the negative strengths to ‘-1’. The reason behind using that discretization step was to only consider the A/B compartment signature irrespective of the compartment strength. Once binarized, the data was then organized in a matrix format, where the rows are the genomic regions and columns are cells and principal component analysis (PCA) was performed on that data. Since we were dealing with a small set of samples (cells) for our analysis, we kept all the principal components from the PCA transformation for further analysis. Then, a k-nearest neighbor graph with 2 nearest neighbors (includes the node itself) was constructed from the PCA transformed data and the network was visualized with a force-directed layout.

### Microarray

RNA was purified from 5 million cells at three different passages, per cell line, using the RNEasy mini kit (Qiagen 74104) using QIAshredder (Qiagen 79654) for homogenization. Purification was followed up by cleanup, and concentration using the RNase-Free DNase Set (Qiagen 79254) and QIAquick PCR Purification Kit (Qiagen 28104), respectively. RNA concentration was determined using a NanoDrop One (Thermo).

Clariom S microarray was carried out through the Transcriptome Analysis Services (Transcriptome Profiling), from Thermo Fisher Scientific.

In addition, for cross-validation purposes, data for HG-U133 Plus2 microarray was also collected from ENCODE, as follows: RWPE (GSM966512, GSM966513, GSM966514), LNCaP (GSM2571978, GSM2571979, GSM2571980), DU145 (GSM1374469), PC3 (GSM1517530, GSM1517531, GSM1517532), LNCaP-C4-2B (GSM1565257. GSM1565258), and 22RV1 (GSM2571966, GSM2571967, GSM2571968).

Analysis of Microarray data was performed using the Transcription Analysis Console (TAC) from Applied Biosystems (Thermo Fisher Scientific), curating the log2 fold upregulated/downregulated genes (p<0.05) with targets identified in the ΔPC1 analysis.

## Results

### Hi-C reveals distinct changes in the 3D genome structure of a cohort of cell lines that model prostate cancer progression

To determine how the genome’s organization is affected by disease stage, we selected a cohort of nine cell lines that model the progression of prostate cancer, as follows: RWPE1 (Bello et al., 1997) was used to represent normal epithelium. LNCaP (Horoszewicz et al., 1983), originally isolated from lymph node metastasis, was used as an early adenocarcinoma model. VCaP (Korenchuk et al., 2001) and MDaPCa2a/b (Navone et al., 1997) were used as models for prototypical osteoblastic bone metastasis; these cells are Caucasian and African American origins, respectively. 22RV1 (Sramkoski et al., 1999) and LNCaP C4-2B (Thalmann et al., 2000) were included in the study as cells that, although isolated from human sources, are models of murine bone metastasis. Atypical metastatic cell lines used in this study include PC3 (Kaighn et al., 1979) (osteoclastic and androgen-independent) and Du145 (Stone et al., 1978) (brain metastasis) (Fig 1).

**Figure 1.**
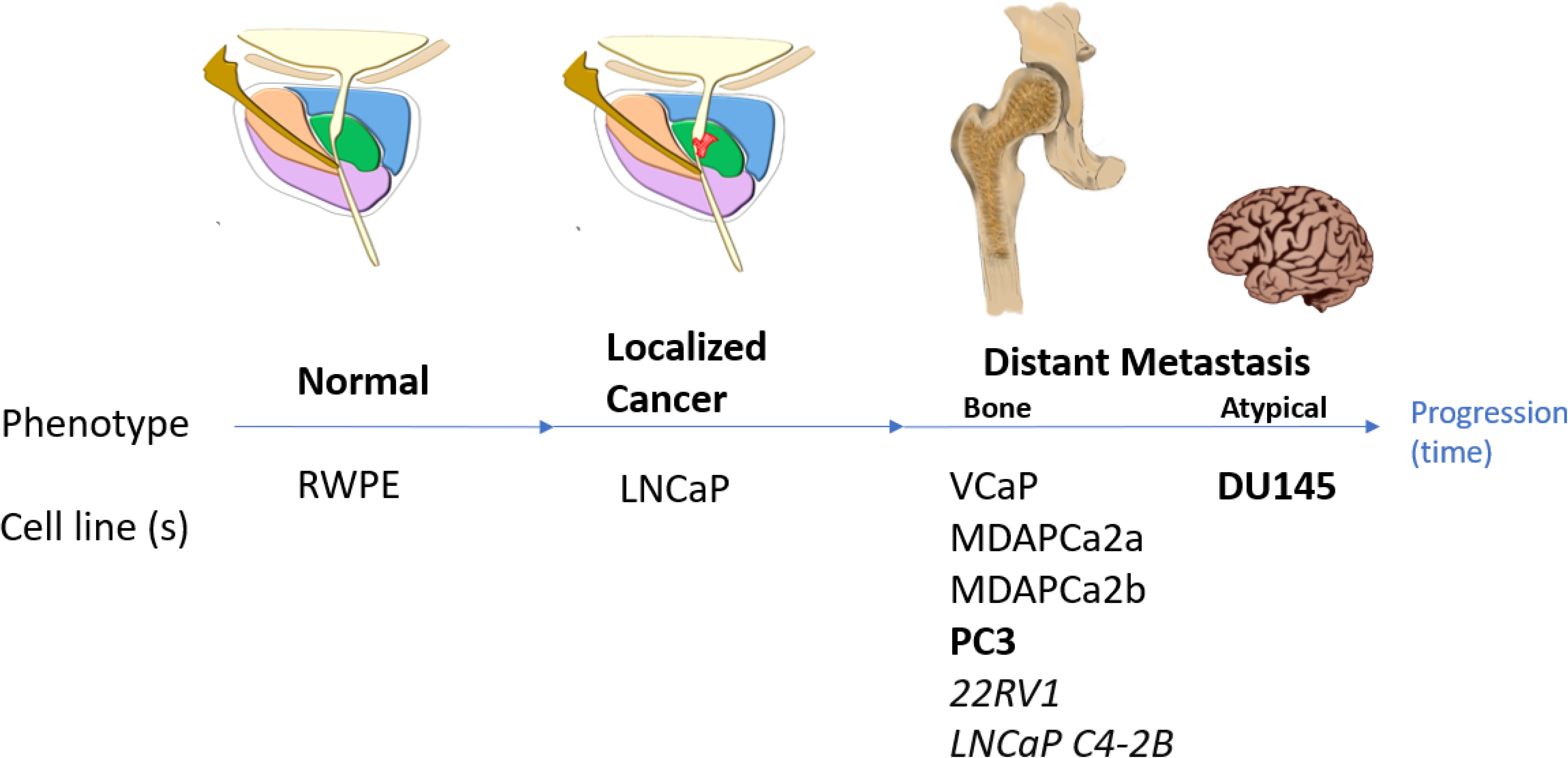
A cell line-based model for the progression of prostate cancer. Cell lines used in this study model different stages in the progression from normal epithelium (RWPE) to localized disease (LNCaP) to prostate cancer that has metastasized to bone (VCaP, MDAPCa2a, MDAPCa2b, PC3) or to an atypical site (DU145). Cell lines developed through mouse xenografts and androgen independent cell lines highlighted in italics and boldface, respectively.

Hi-C was performed in cells under normal cell culture conditions (Methods) on five million cell pellets. Additionally, publicly available datasets for RWPE and LNCaP-C42B were used as comparisons (Rhie et al., 2019, Taberlay et al., 2016). Hi-C mapping and quality control statistics for all samples can be found in Sup. Fig. 1.

The frequency of chromosome contacts is represented in a heatmap where the XY axes are the chromosomal coordinates. The color intensity reflects the frequency with which two particular locations were found to be in contact. At a resolution of 1 megabase (Mb) bins, the Hi-C heatmaps reveal chromosome territories. At a 250-kilobase (kb) resolution, it is possible to see a plaid pattern of interaction strength, which represents the spatial segregation of A and B compartments. We classify each genomic region as belonging to the A or B compartment using principal component analysis. Positive values of the first eigenvector (eigen1) represent A compartment regions (typically open euchromatin) while negative values denote B compartment (heterochromatin). Finally, at 40 kb resolution, distinct topologically associating domains (TADs) are evident, which are regions of enhanced contacts (which may promote contacts between promoters and enhancers) segregated by the insulator protein CTCF (Fig. 2A).

**Figure 2.**
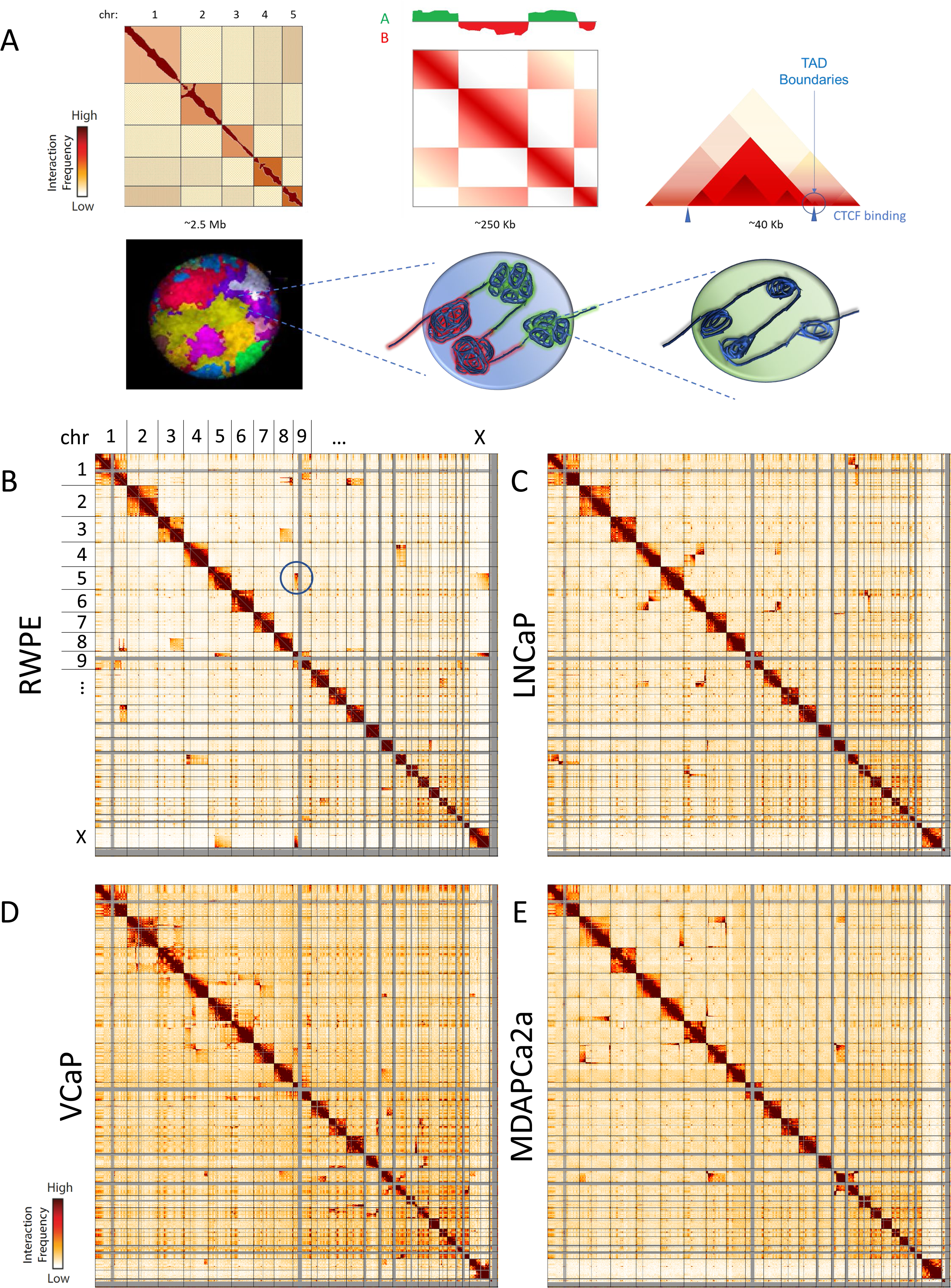
Chromosome conformation capture characterizes the hierarchical genome of each cell line. A) At 1 Mb resolution (left), Hi-C heatmaps reveal show defined chromosome territories along the diagonal. At 100 kb resolution a characteristic plaid pattern emerges: principal component analysis of this matrix reveals A/B chromosome compartmentalization (center). At 40 kb resolution, topologically associating domains (TADs), within compartments, become apparent (right). 2.5 Mb Hi-C heatmaps for RWPE (B), LNCaP (C), VCaP (D) and MDAPCa2a (E). Translocations between chromosomes appear as high interaction frequency areas away from the diagonal. Translocation between chromosomes 8 and 5 highlighted as an example in RWPE (circle). While these translocation events have been validated by SKY analysis, the high resolution of Hi-C data allows for the characterization of smaller events (Sup. Fig. 2).

Whole-genome contact maps for cells in our model reveal that the highest incidence of interactions occurs in cis: chromosomes primarily interacting within themselves (Fig 2B, C, D, E). Chromosomal translocations are evident as very strong interactions occurring in trans between different chromosomes. Comparing our Hi-C results with published spectral karyotyping (SKY) data (Pan et al., 1999, van Bokhoven et al., 2003) and karyotyping information from ATCC, we found that our Hi-C data detects 84% of all previously reported translocations. Owing to the high resolution of Hi-C data, we also characterized several previously unreported, smaller translocation events (Sup. Fig. 2). Interestingly, chromosomal territories remain well defined throughout progression, with subtle changes in intrachromosomal interactions, as shown in 250 Kb-resolution heatmaps of each chromosome (Sup. Fig. 3).

### Genes important to prostate cancer progression switch chromatin compartment identity, and these changes are accompanied by transcription activation

Examination of higher resolution (250 kb) z-score correlation heatmaps of cis interactions for each chromosome shows that the characteristic plaid contact pattern, associated with spatial compartment identity, changes among cell lines. For example, for chromosome X, an erosion of the pattern is observed in the bone metastatic cell line VCaP compared to the normal epithelium (RWPE). In contrast, the patterning observed in the model cell line for adenocarcinoma (LNCaP) and the bone metastatic MDAPCa2a line is more similar for chrX (Fig.3A). These patterns can be observed throughout all chromosomes, and the changes in compartmentalization are specific to each cell line (Sup. Fig.4). Motivated by the visible compartment “erosion” in VCaP, we quantified the overall strength of interactions within A compartment regions and within B compartment regions in the different cell types. We find that all prostate cancer cell lines show a loss of A compartment interaction strength relative to RWPE, reflecting an increased intermixing of the A compartment with B compartment regions. Meanwhile, B compartment strength experiences less change in LNCaP and VcaP and even increases in MDAPCa cell lines. Overall, this leads to an imbalance of B and A compartment strength in all prostate cancer cell lines (Sup. Fig. 5A).

**Figure 3.**
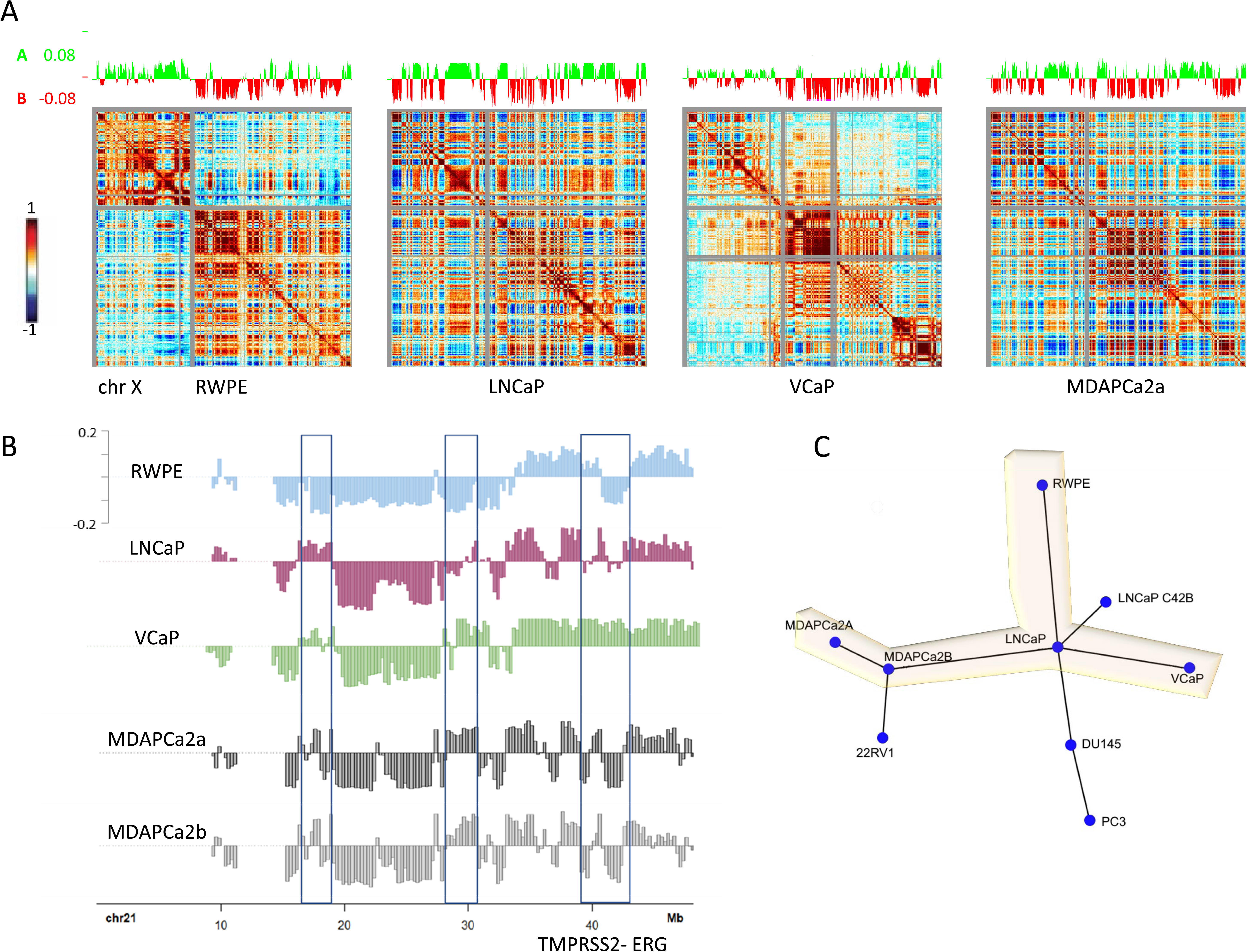
Changes in A/B compartment identity are relevant to prostate cancer progression. A) 250 kb resolution z-score correlation heatmaps of cis interactions along the X chromosome show a characteristic plaid pattern that denotes compartment identity (compartments classified by PC1 of this matrix: green = A, red = B). A weakening of this pattern is observed in the bone metastatic cell line VCaP when compared to normal epithelium (RWPE). Similar patterning is observed in the model cell line for adenocarcinoma (LNCaP) and the bone metastatic MDAPCa2a line. B) PC1 compartment tracks for chromosome 21 show distinct hotspots of compartment identity switches (boxes) along the chromosome between the normal epithelium (RWPE) and bone metastatic cell lines. The right-most region encompasses the TMPRSS2-ERG locus. C) A comparison of genome-wide compartment tracks for all cell lines by nearest neighbor analysis reveals that LNCaP is a central node in a model axis to metastasis (highlighted yellow), connecting RWPE (normal epithelium) to bone metastatic cell lines (VCaP and MDA cell lines). Atypical androgen independent lines cluster together along a different axis from LNCaP.

Systematic analysis of the A/B compartment tracks, per chromosome, for all the cell lines in our model, revealed regions where the compartment identity remained the same and where it was changed (Sup. Fig. 5B and 6). From the contact maps, it is evident that some chromosomes in some cell lines are broken into multiple pieces (Sup. Fig. 7). However, when we perform compartment analysis on each of these broken pieces separately, we find that their underlying A/B compartmentalization is largely similar to cell lines with unbroken chromosomes. We note that there is a general level of similarity between epithelial-derived cell lines (Sup. Fig. 5B). By comparing compartment tracks for each cancer cell line to the normal control (RWPE), we find that the changes among cell lines are specific and localized. For example, eigenvector tracks for chromosome 21 show three distinct hotspots of compartment identity switches between the RWPE and cancer cell lines (Fig 3B). Interestingly, the right-most region encompasses the TMPRSS2-ERG locus, a local translocation site that correlates with worse progression and metastasis in prostate cancer (Tomlins et al., 2005, Demichelis et al., 2007, Hagglof et al., 2014).

A comparison of genome-wide compartment tracks for all cell lines by nearest neighbor analysis (SPRING Plot, see Methods) revealed that LNCaP is a central node in a model axis to metastasis (Fig. 3C, axis highlighted in yellow), connecting RWPE (normal epithelium) to bone metastatic cell lines (VCaP and MDAPCa). Interestingly, the compartment pattern of MDAPCa2A and 2B, which were derived from an African American patient, is distinct from VCaP, which is of Caucasian lineage. This analysis also clusters atypical metastatic lines DU145 and PC3 together along a third axis radiating from the LNCaP central node.

To consistently mathematically classify regions that significantly changed compartments between cell lines, PC1 values per bin per cell line were normalized by subtracting the corresponding value derived from the normal RWPE cell line. Significant changes in compartment identity (|*X̅* + 1.5*σ*|) were identified, and the corresponding genomic areas annotated (Sup. Fig. 8). Through this analysis, we have identified 181 genomic bins (250 kb in size) whose compartmentalization changes from the B to the A compartment in either (a) six or more cancer cell lines compared to RWPE or (b) in all cells in the progression axis, compared to RWPE. These genomic bins contain about three hundred genes that are therefore moving from the B to the A compartment (Sup. Table 2), suggesting that these genes become more likely to be transcribed. These include androgen receptor (AR), WNT5A, CDK14, and genes located close to TMPRSS2, such as BACE2. The majority of these genes (256) are grouped in forty-eight proximal clusters (Sup. Table 3). We have also identified eighty-six genes whose genomic loci switch compartments from the A to B compartment, suggesting a genome structure rearrangement more likely to result in gene silencing (Sup. Table 2). Such is the case for certain cadherins, annexins, and mediators of inflammation. Of these genes, sixty-four are grouped in sixteen clusters (Sup. Table 3).

Since a switch from the B to A compartment could signify transcriptional activation, we used the Clariom-S microarray to profile expression levels in the targets identified. Using the expression level of RWPE as a baseline, we have found that forty-seven percent of genes that switch from B to A compartment show a significant transcriptional induction in LNCaP (2-fold or higher). In turn, forty-nine percent of those genes are even further upregulated in VCaP compared to LNCaP, suggesting an exacerbation of this expression pattern with progression. These genes include, among others, androgen receptor (AR), TMPRSS2, CDK14, and WNT5a. (Fig. 4A). In contrast, only 18% of the genes that show induction in LNCaP are induced further in the MDAPCa cell lines. (Fig. 4B).

**Figure 4.**
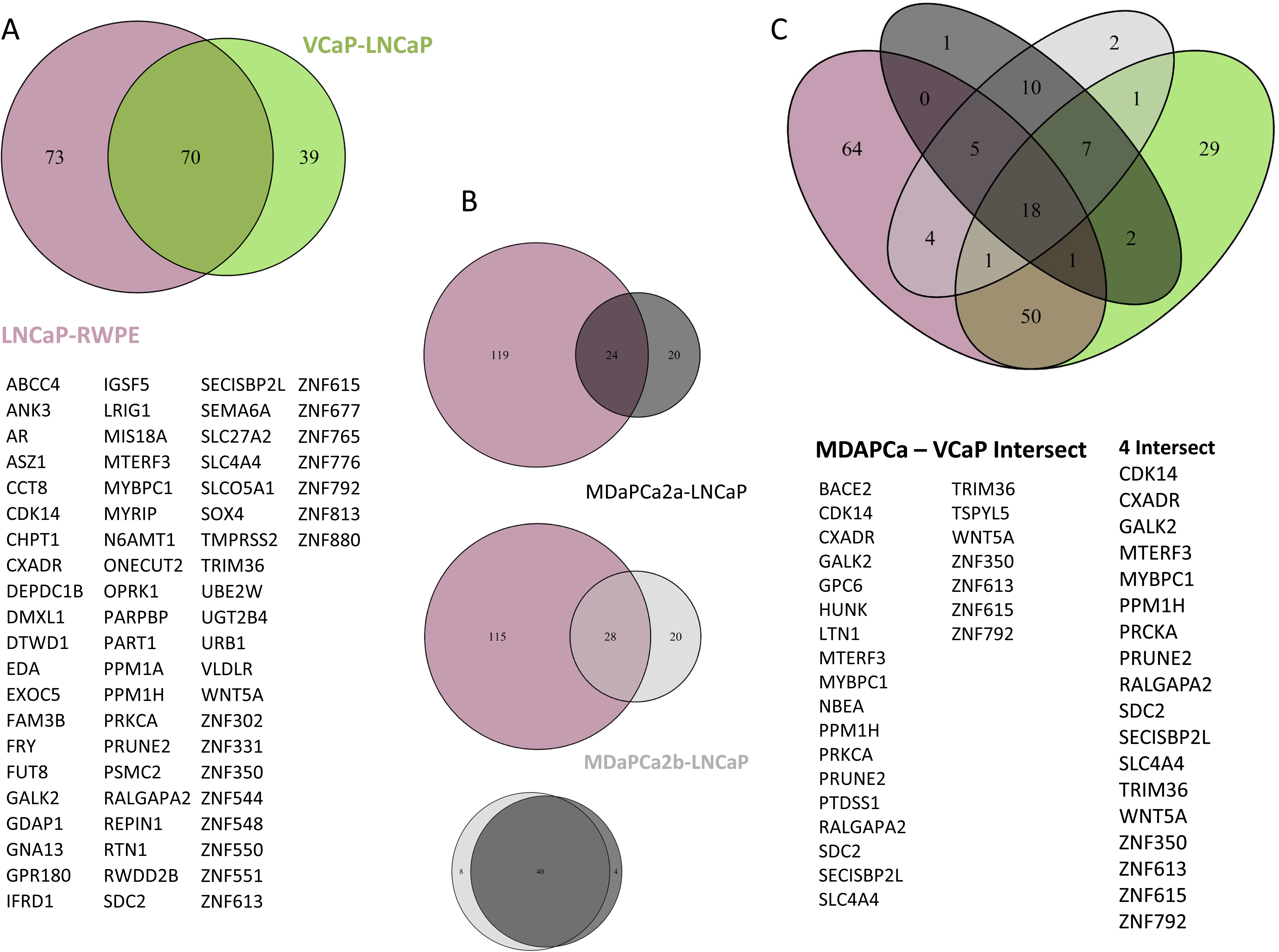
Microarray: genes whose increased expression correlates with changes in B to A compartment identity. A) 143 genes identified as switching from the B to the A compartment show overexpression in LNCaP when compared to RWPE (purple). Of these, 70 are further overexpressed in VCaP (green) compared to LNCaP, denoting an exacerbation of this expression pattern with progression. This overlapping gene set includes known prostate cancer targets such as androgen receptor (AR) and TMPRSS2. B) Of the genes upregulated in LNCaP vs. RWPE (purple) 24 and 28 genes are further overexpressed in MDAPCa2a and MDAPCa2B, respectively, when compared to LNCaP. Overlap of the genes overexpressed in the MDA cell lines, compared to LNCaP, shows a high concordance between these cell lines, which were derived from the same patient. C) Genes that change compartment identity from B to A and are overexpressed in all bone metastatic cell lines, compared to LNCaP, which in turn are overexpressed in LNCaP compared to normal epithelium.

Finally, we found that 25 genes that underwent a B to A compartment switch show transcriptional induction in both MDAPCa cell lines relative to VCaP, including CDK14, WNT5a, and BACE2, which is a close neighbor of TMPRSS2. (Fig 4C). This data suggests that these transcriptional hotspots are common for osteoblastic metastatic cells and might be necessary for colonization and survival in a secondary bone site.

### Compartment identity switches are accompanied by distinct structural changes at the TAD level, including boundary shifts

Topologically associating domains (TADs) contribute to the 3D architecture of the genome by sequestering enhancers and promoters with their target genes, with a low likelihood of interaction across boundaries (Lupianez et al., 2015, Nora et al., 2012, Dixon et al., 2012). The disappearance of TAD boundaries has been identified as a potential activator of oncogenes (Hnisz et al., 2016). To analyze whether there are changes in 3D genomic structure at the local level surrounding genes associated with compartment identity switches, we used higher resolution Hi-C heatmaps (40Kb).

The WNT5a-ERC2 locus (Fig. 5 Cluster 1) is in the B compartment in normal epithelium, switching to the A compartment in all four members of the model axis to progression: LNCaP, VCaP, and both MDAPCa cell lines. While the right TAD boundary location is relatively consistent, the location of the left boundary shifts positions: In LNCaP, DU145, MDAPCa2B, and PC3, WNT5A localizes in a TAD with LRTM1 instead of ERC2. Interestingly, in VCaP, these three genes are clustered in the same TAD. FZD1 and CDK14, a receptor and activator cyclin of non-canonical WNT signaling, and which have been associated with the function of WNT5A, also switch to the A compartment (Fig. 5, Cluster 2). In this case, the genes are separated by a TAD boundary that rests atop the gene body of CDK14, but increased loops are evident on the separate TADs in the cancer cell lines, compared to normal epithelium. Remarkably, CDK14 is one of the genes whose transcription is consistently upregulated in all metastatic cell lines in the progression axis. TAD boundary shifting, appearance and disappearance were also observed in other clusters that switch compartment identity (Examples in Sup. Fig 9).

**Figure 5.**
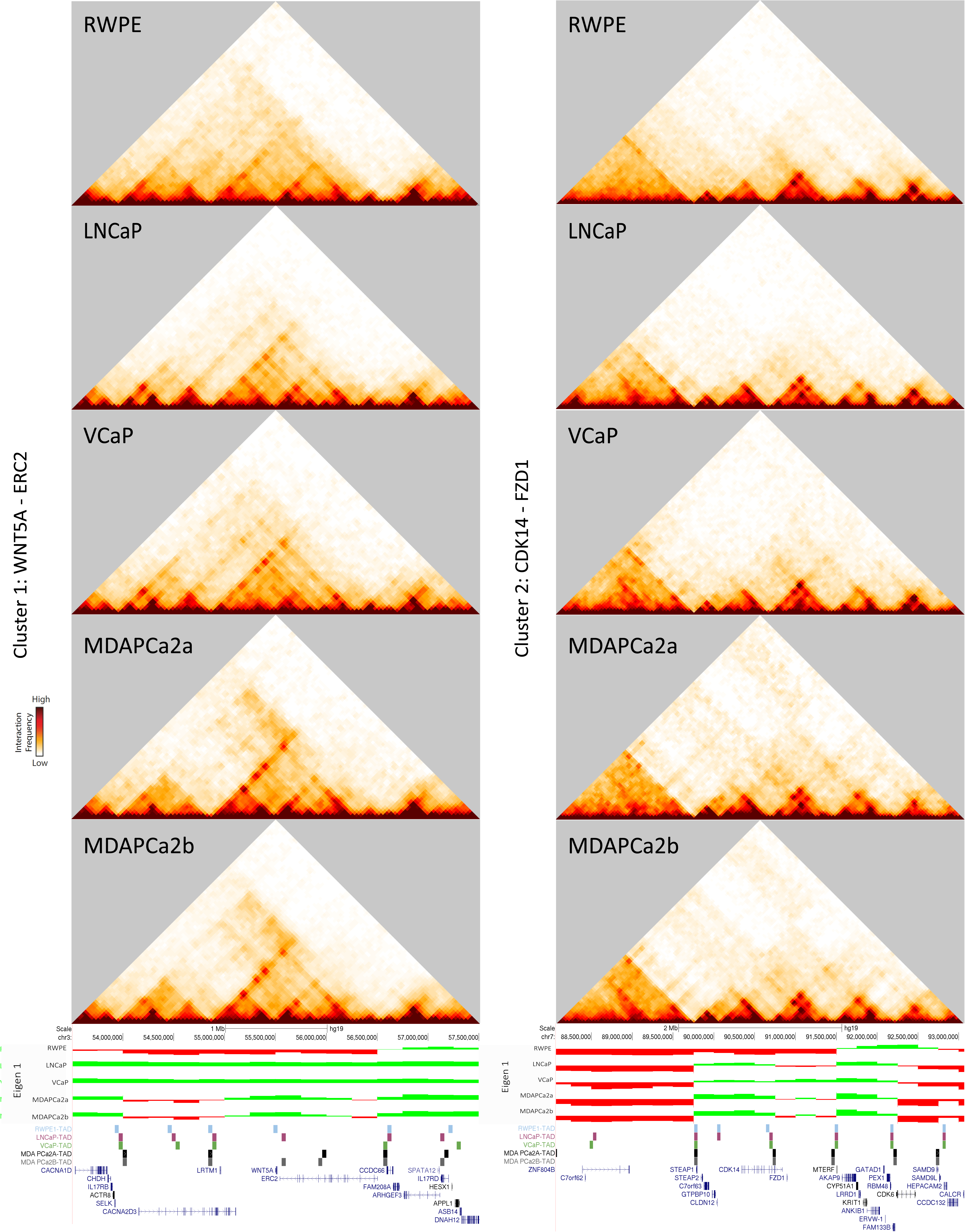
Gene clusters that switch from the B to the A compartment include genes critical for prostate cancer progression. 40 kb resolution heatmaps, compartment tracks, and TAD boundaries are shown for two representative clusters of genes that change compartment across the primary axis of progression. (Left) WNT5A switches from the B to the A compartment in cells of the primary progression axis. WNT5A sits at the location of a shifting TAD boundary, and strong interaction sites in the TAD surrounding this gene become more apparent in cancer progression as compared to RWPE. (Right) CDK14 switches from the B to the A compartment in most cancer cell lines. A TAD boundary localizes atop the gene body of CDK1.

A detailed survey of all compartment-switched loci revealed five possible categories of changes in local structure. We classified all compartment switch regions into these categories. Strikingly, the androgen receptor locus, displays all five types of structure change across the different cell lines. First, a highly disorganized area (No TADs, uniformly distributed interactions across a region of the heatmap) becomes highly organized, or vice versa (Fig 6A). In AR, the gene is in the B compartment in RWPE, and the area around it is highly disorganized. With progression, the area becomes organized into TADs and sub-TADs for both LNCaP C4-2b and VCaP. The third type of structural conformation is the appearance of a high incidence of interactions along the edge the TAD (Fig. 6B). We call these interactions “loops” or “stalled loops” because these have previously been associated with the phenomenon of cohesin becoming stalled as it extrudes loops but encounters RNA polymerase, CTCF, or other barriers. This increased loop formation is evident at the AR locus in LNCaP, MDAPCa2a, and MDAPCa2b. The fourth structural feature is a complete absence of structure (loss of contacts in the entire region) associated with the gene of interest. This “structural desert” is observed at the AR locus in the 22RV1 cell line (Fig. 6C) and is reminiscent of previously observed structural features at highly transcribed long genes(Leidescher et al., 2020, Heinz et al., 2018). Finally, the local structure can remain unchanged. For AR, this is the case in PC3 and DU145, where the gene remains in a disorganized region of the B compartment (Fig. 6D).

**Figure 6.**
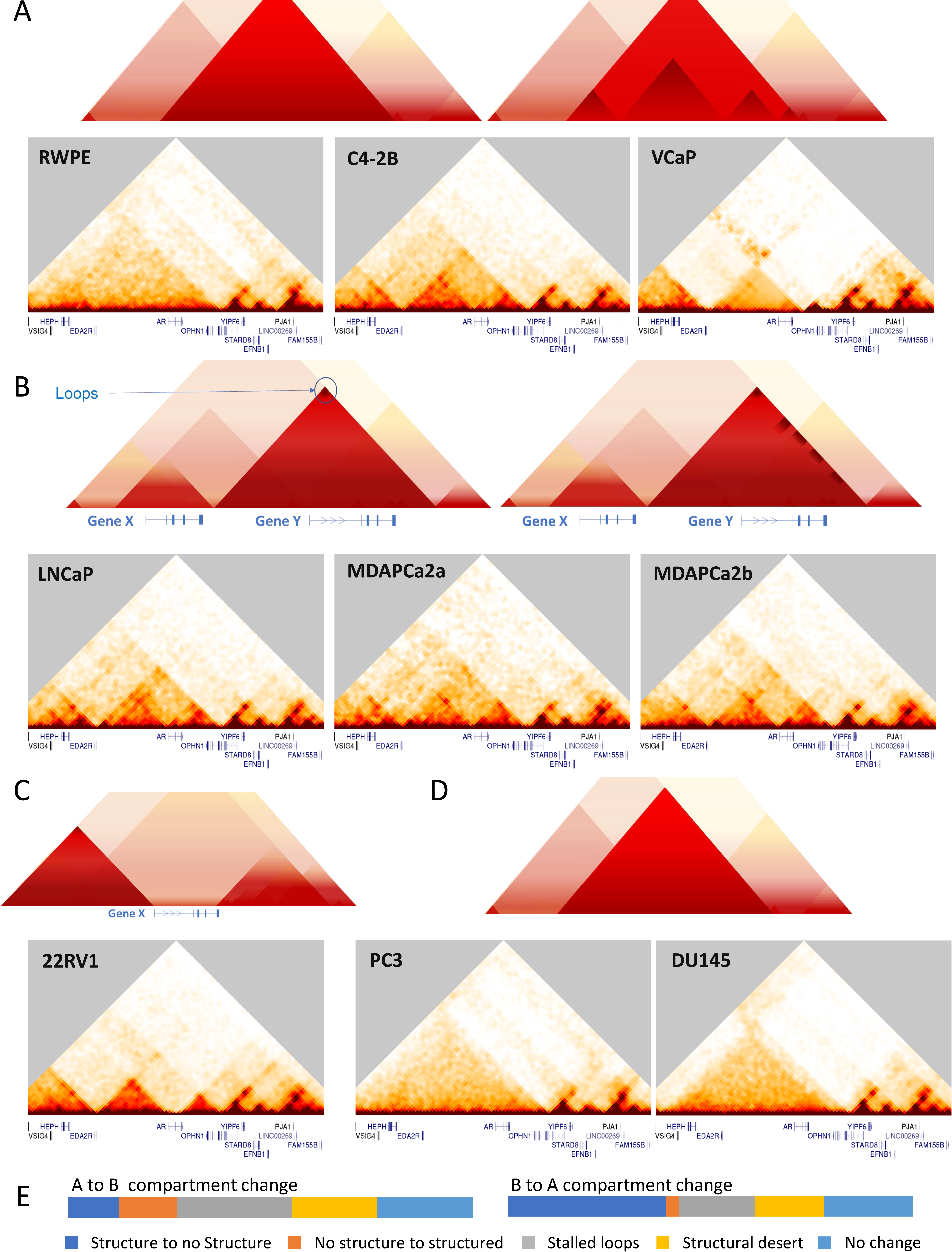
Compartment identity switches are accompanied by varying types of structural changes at the TAD level. The AR locus is shown as an example of the different categories of TAD-scale genome organization changes that accompany compartment changes. A cartoon of the general type of change is shown above sets of representative data. A) No structure to structure. In RWPE, the AR locus is in the B compartment, and there is little defined TAD structure. TADs that appear C4-2B and VCaP are of different sizes and location. B) Stalled loops. In addition to the appearance of a TAD structure, LNCaP, MDAPCa2a and MDAPCA2b show enriched foci of interactions along the edges of TAD boundaries, consistent with previously documented cases of stalled loop extrusion. C) Structural desert. In 22RV1, the entirety of the AR gene is in a region of depleted contacts. D) No change. In PC3 and DU145, the AR locus remains disorganized as in RWPE. E) Genome-wide distribution of above structural change categories across regions that change from A to B (left) or B to A (right).

Overall, for those genes that switch from the A to the B compartment, all types of changes happen with fairly even probability: 12.64% lose structure at the local level, 14.37% gain structure, 28.16% present stalled loops, 21.26% associate with structural deserts, and 23.56% do not change. However, genes located in the B to A compartment switches predominantly acquire structure (39.06%) or do not change (21.88%). Only about 3% of B to A switch loci lose structure. The remaining loci are distributed evenly between stalled loop-areas and structural desert change (Fig 6E). From this survey of prostate cancer cell lines, therefore, we also gain basic insight about the types of local structure change that most often accompany compartment level changes.

### The TMPRSS2-ERG locus shows an increase in local interactions in cell lines in the metastatic progression axis

As previously mentioned, the incidence of the TMPRS2-ERG translocation has a positive correlation with prostate cancer progression and a poor prognosis. For the TMPRSS2-ERG locus, we observe that the region that encompasses TMPRSS2, BACE2, PLAC4, MX1, MX2, and FAM3B switches from the B compartment in RWPE to the A compartment in all cancer cell lines. In the model cell line for normal epithelium RWPE, this genomic area is enclosed in a large TAD, including genes downstream of TMPRSS2 (Fig. 7A), as is also the case for LNCaP, 22RV1, MDAPCa2a, and PC3, with a slight shifting of the TAD boundary location. In LNCaP-C42B, an osteoblastic cell line derived from LNCaP, a sub-TAD appears within this locus, sequestering the MX1 gene. Sub-TAD fragmentation also occurs in DU145, MDAPCa2b, and VCaP. In contrast, ERG is in the A compartment in RWPE (normal epithelium), switching to the B compartment only in LNCaP and MDAPCa2b. The TAD boundary location around ERG does not change. Of note, the whole genomic area surrounding these two clusters is located in the A compartment in VCaP, a known carrier of the TMPRSS2-ERG translocation, evident as a high interaction location in the Hi-C heatmap (Fig 7A: VCaP-rectangle).

**Figure 7.**
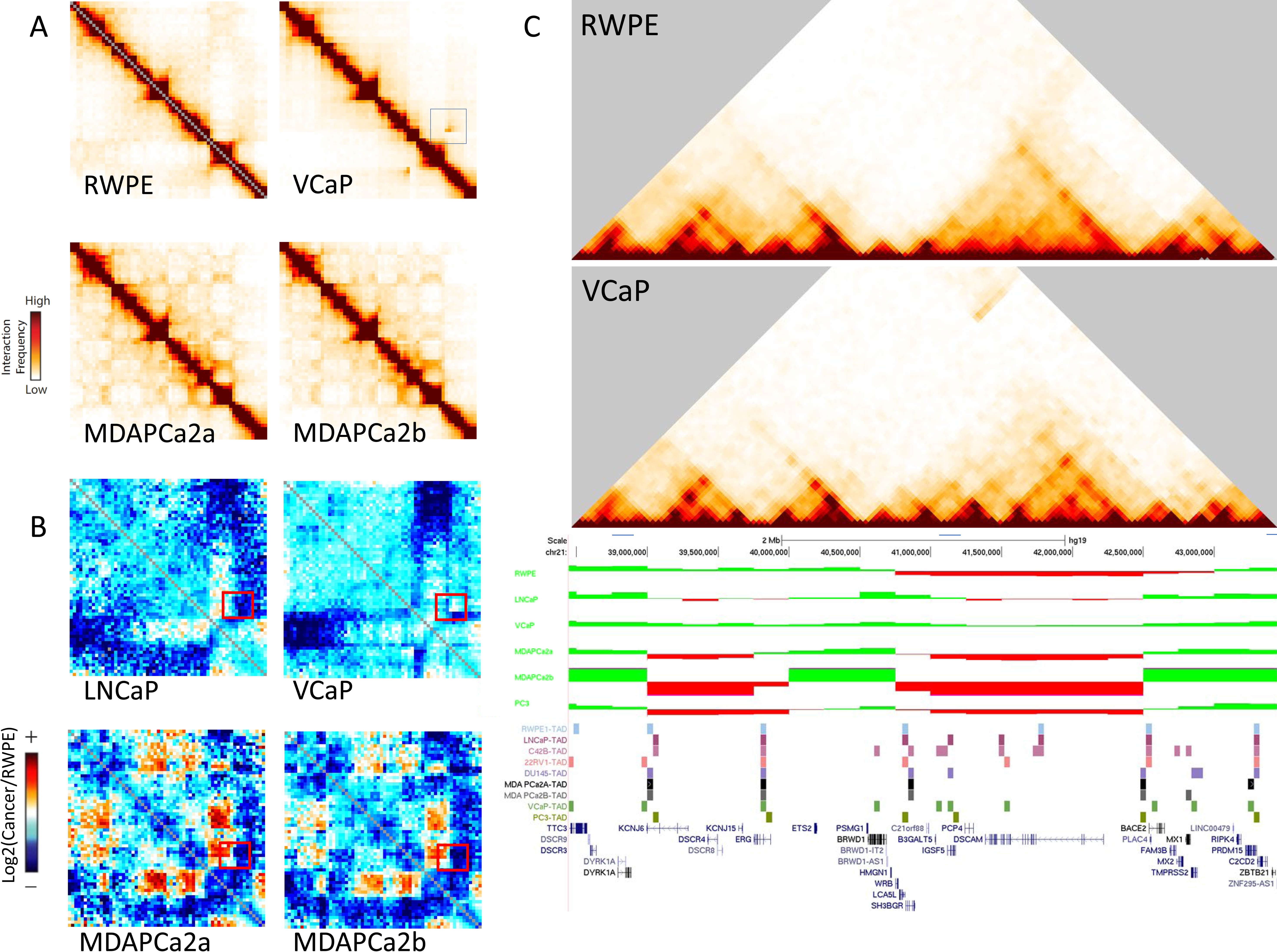
Alterations in interactions around the TMPRSS2-ERG locus in cell lines in the metastatic progression axis. A) Interaction heatmaps of the portion of chr21 containing TMPRSS2-ERG: chr21:30 Mb – 45.2 Mb at 250kb resolution. The TMPRSS2-ERG translocation is visible in VCaP (blue square) as a high interaction away from the diagonal. Interestingly, the other bone metastatic cell lines (MDAPCa2a and b) also show enhanced interactions close to that area. B) Log2 ratio of interactions in each cell line compared to RWPE. Red areas have more interactions in the cell line compared to RWPE, while blue represents fewer interactions compared to RWPE. The area adjacent to the TMPRSS2-ERG translocation (red box) shows an increase in interactions in LNCaP and MDA cell lines. This increase of interactions in the region surrounding these genes is no longer evident in VCaP, where the TMPRSS2-ERG translocation has occurred. C) TMPRSS2 switches from the B to the A compartment in all cancer cell lines while ERG switches from A to B only on certain cell lines. The TMPRSS2-ERG translocation is easily visible in VCaP, as a high interaction area away from the diagonal. Changes in TAD boundary locations around TMPRSS2 are also observed.

In the bone metastatic cell line VCaP the TMPRSS2-ERG translocation is apparent in the Hi-C heatmap of the long arm of chromosome 21 at a 250kb resolution (Figure 7B). While a translocation is not evident in the MDAPCa cell lines, there is a distinct higher incidence of interactions close to that area: a log2 ratio comparison against normal epithelium (RWPE) shows that that chromosomal region is enriched for interactions in adenocarcinoma (LNCaP) (Fig 7C), and that this phenotype is aggravated in both bone metastatic MDAPCa cell lines.

## Discussion

The deliberate, hierarchical organization of chromatin within the eukaryotic nucleus’s constrained space is necessary for adequate DNA maintenance, repair, and gene transcription or silencing, all of which contribute to the cell’s homeostasis. In this study, we have characterized the genomic architecture in a cohort of cells that model prostate cancer progression. At the genome-wide level, given the inherent high resolution of Hi-C data, we have identified small translocations that, to our knowledge, have not previously been reported in the literature.

Many regions across the genome are unchanged in their spatial compartmentalization across cell lines. This suggests that there are inherent 3D genome structure features of prostate epithelium that arise during initial differentiation and tissue patterning (Flyamer et al., 2017, Ke et al., 2017), and that these are persistent, regardless of malignancy status. These results suggest that genomic loci that switch compartment identity between the normal epithelium and cancer cells are associated with an oncogenic genomic architecture profile and that those features result from concerted biological events.

It is noteworthy that the majority of the compartment changes we identify involve a switch from the B to the A compartment. Correspondingly, we find in our compartment strength analyses that the A compartment becomes more intermixed, interacting more broadly, while the strongest B compartment regions remain more spatially segregated in the cancer cell lines. Both of these results point to a general shift in the prostate cancer lines toward a more open / poised for activation chromatin environment, which could lead to misactivation of oncogenes.

Our genome-wide compartment analysis revealed that compartment identity alone is enough to stratify prostate epithelium in a continuum of progression. LNCaP is a central node that connects the normal epithelium (RWPE) to bone metastatic cell lines (MDAPCa2a/b and VCaP). These results provide further insight into the importance of cell line selection in progression studies: the atypical metastatic cell lines DU145 and PC3 cluster together away from the primary axis. This emphasizes that as researchers select cell lines for study, it is important to consider the differences in these cell lines and that not all will capture the most common pathways of metastasis.

It is important to consider that the compartment switching events observed here involve clusters of genes, that in some cases expand through several topologically associating domains (TADs). This observation echoes previous results in a plant system, where clusters of genes often changed their spatial compartmentalization and expression together (Nutzmann et al., 2020). Interestingly, the clusters we observe include both genes previously identified in prostate cancer progression and others seemingly unrelated. Such is the case of the WNT5a locus in which is a known target of both prostate tissue development, patterning, and cancer aggressiveness (Allgeier et al., 2008, Dai et al., 2008, Yamamoto et al., 2010, Huang et al., 2009). Our results show that in normal epithelium, WNT5a is sequestered within a TAD with the ELKS/RAB6-Interacting/CAST Family Member 2 (ERC2). In contrast, TAD boundary shifting or eviction in the proximity of WNT5a in the bone metastatic cell lines represented in our primary axis for progression (VCaP and MDAPCa2a/b) results in TAD-limited interactions with Leucine-rich repeats and transmembrane domains-containing protein 1 (LRTM1) instead. While this shift does not result in transcriptional induction of LRTM1, it could potentially lead to aberrant interactions between the promoters for both genes, ultimately resulting in the overexpression of WNT5a. Since TAD structure is essential for proper gene regulation (Lupianez et al., 2015, Rhie et al., 2019, Guo et al., 2018), this phenomenon requires further exploration. Still, it is attractive to consider TAD-targeted therapies that hold the potential to reverse the deleterious effects of TAD shifting. Interestingly, another target of non-canonical WNT signaling, CDK14 (reviewed by Davidson and Niehrs, 2010), also switches from the B to the A compartment and is transcriptionally activated in our metastatic axis.

The clustering of different genes in the described compartment switches raises the interesting question of whether certain genes could act as a driver of compartment identity switches while neighboring genes act as “passengers.” For example, the transcriptional activation of one gene could influence a whole genomic region to switch to the A compartment. Indeed, previous work has shown that binding of transcriptional activators can prefigure spatial compartment alterations (Stadhouders et al., 2018, Therizols et al., 2014). This compartment switch would then result in the switching of neighboring genes, which may increase their probability of later becoming activated as well. Recent work has shown that spatial reorganization of a chromosome region can make it more permissive for derepression, even if the structural switch does not immediately change its expression level (Manjón et al., 2021). Is it possible that earlier transcriptional events required for the cell to survive a particular insult trigger a full compartment shift? Is this, in turn, a potential trigger for oncogenic transcriptional activity?

Such seems to be the case of the observed amyloid precursor protein (APP) – beta secretase 2 (BACE2) axis, along chromosome 21. APP presents with enhanced expression in the LNCaP – MDAPCa – VCaP axis, compared to the normal RWPE (Sup. Fig 10). APP is also proximal to one of the persistent compartment switches in chromosome 21, but it does not change compartment itself: The A compartment identity atop the gene gets stronger with progression. Mounting evidence from the Alzheimer’s field, where abnormal amyloid processing results in aggregation and neurodegeneration, has shown that this protein and derived peptides serve a crucial role as antimicrobials and are necessary for mounting an appropriate host response to infection (thoroughly reviewed by Moir et al., 2018). If an environmental signal such as infection results in the prostate epithelium being exposed to excessive or chronic APP, could this trigger expression of its processing enzyme BACE2? Our evidence of BACE2 switched compartmentalization and experienced higher levels of transcription in all three prototypical bone metastatic lines suggests so. Critically, these events would imply the need for the compartment switch around the TMPRSS2 locus.

Since its discovery, the TMPRSS2-ERG translocation in chromosome 21 has been recognized as an important indicator of poor prognosis and a higher risk of prostate cancer-related death (Tomlins et al., 2005, Perner et al., 2006, Demichelis et al., 2007, Hagglof et al., 2014, Deplus et al., 2017). The genesis of this translocation and fusion, however, remains poorly understood. Here, we implicate a 3D genome organization change of compartmentalization in this event. As mentioned before, TMPRSS2 switches from the B to the A compartment consistently in all prostate cancer cell lines we queried. Meanwhile, ERG switches from the A to the B compartment in LNCaP and its nearest neighbor in our experimental metastatic axis, MDAPCa2b. Remarkably, ERG remains in the A compartment for VCaP and MDAPCa2a, suggesting a potential dual switching event during progression to more aggressive phenotypes. All these spatial rearrangements in such proximity result in the enhanced contact incidence observed in all of these cell lines, even the early adenocarcinoma model LNCaP (as described in figure 7), and ultimately could lead to the translocation event, as it has been shown in other systems (McCord and Balajee, 2018, Zhang et al., 2012)

Recent efforts (Hawley et al., 2021), have characterized 3D genomic profiles in prostate tumor cohorts. These studies recapitulate our findings that the 3D genome organization between malignant and benign prostate tissues remains largely consistent. We propose that prostate cancer progression is associated with specific changes in the 3D genome structure that arise early in the disease and facilitate an oncogenic expression phenotype. Based on these results, we can hypothesize that analyzing the 3D genome structure of patient derived samples could be a prognostic marker for progression and bone metastasis.

## Acknowledgments

We thank Jeremy Hughes (Web Communications Manager – UTK Department of Arts & Sciences) for his help in constructing the accompanying website.

We thank Nora Navone Ph.D. for providing the MDAPCa cell lines. We thank Justin Roberts Ph.D. for technical support in the culturing of these cell lines.

We thank Rosela Golloshi Ph.D. and Jacob Sanders Ph.D. for their technical support. This work was supported, in part, by the by NIH NIGMS grant R35GM133557 to R. P. McCord. R. San Martin was supported by a Postdoctoral Fellowship from the American Cancer Society (134060-PF-19-183-01-CSM). The authors declare no competing financial interests.

## Data Availability

All Hi-C and microarray data are available on GEO at accession number GSE172099. Other processed data figures, and a UCSC Genome Browser Track Hub containing all data are available for browsing at https://3dgenome.utk.edu/3d-genome-architecture-in-prostate-cancer-progression/.

## Author Contributions

*Conceptualization:* R. San Martin and R. P. McCord; *Investigation:* R. San Martin. *Formal analysis:* R. San Martin, R. P. McCord P. Das and R. Dos Reis Marques; *Visualization*: R. San Martin, Y. Xu and R. Dos Reis Marques; *Writing – original draft*: R. San Martin and R. P. McCord; *Software*: P. Das and Y. Xu *Writing – review and editing*: all authors; *Supervision*, R. P. McCord.

## Supplementary figures

**Note:**

The following figures are presented in an abridged form:

Supplementary figure 1

Supplementary figure 2

Supplementary figure 3

Supplementary figure 4

Supplementary figure 6

Please visit https://3dgenome.utk.edu/3d-genome-architecture-inprostate-cancer-progression/ to access the complete supplementary figures.

**Supplementary Figure 1.**
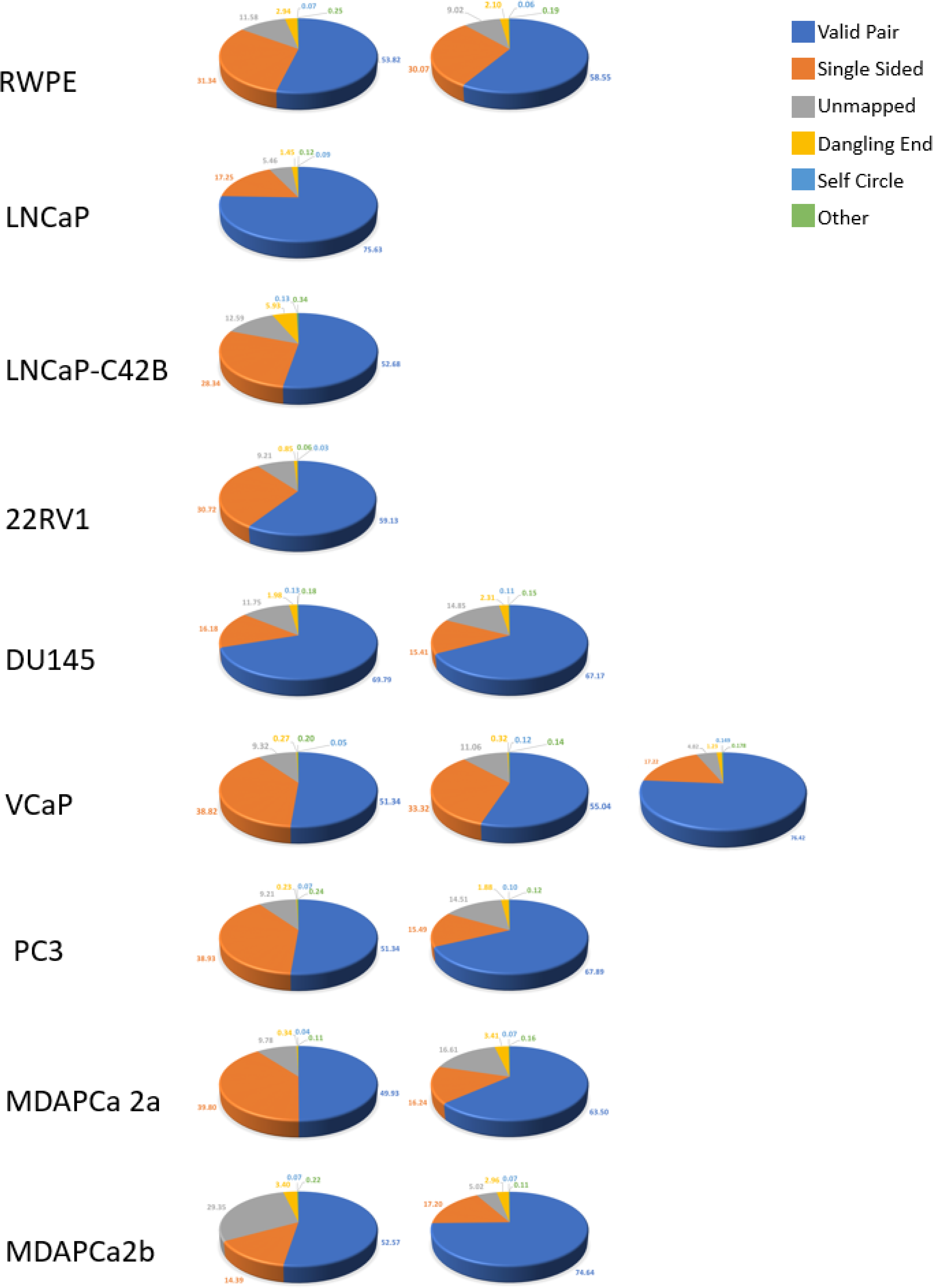
Mapping Statistics for all Hi-C samples.

**Supplementary Figure 2.**
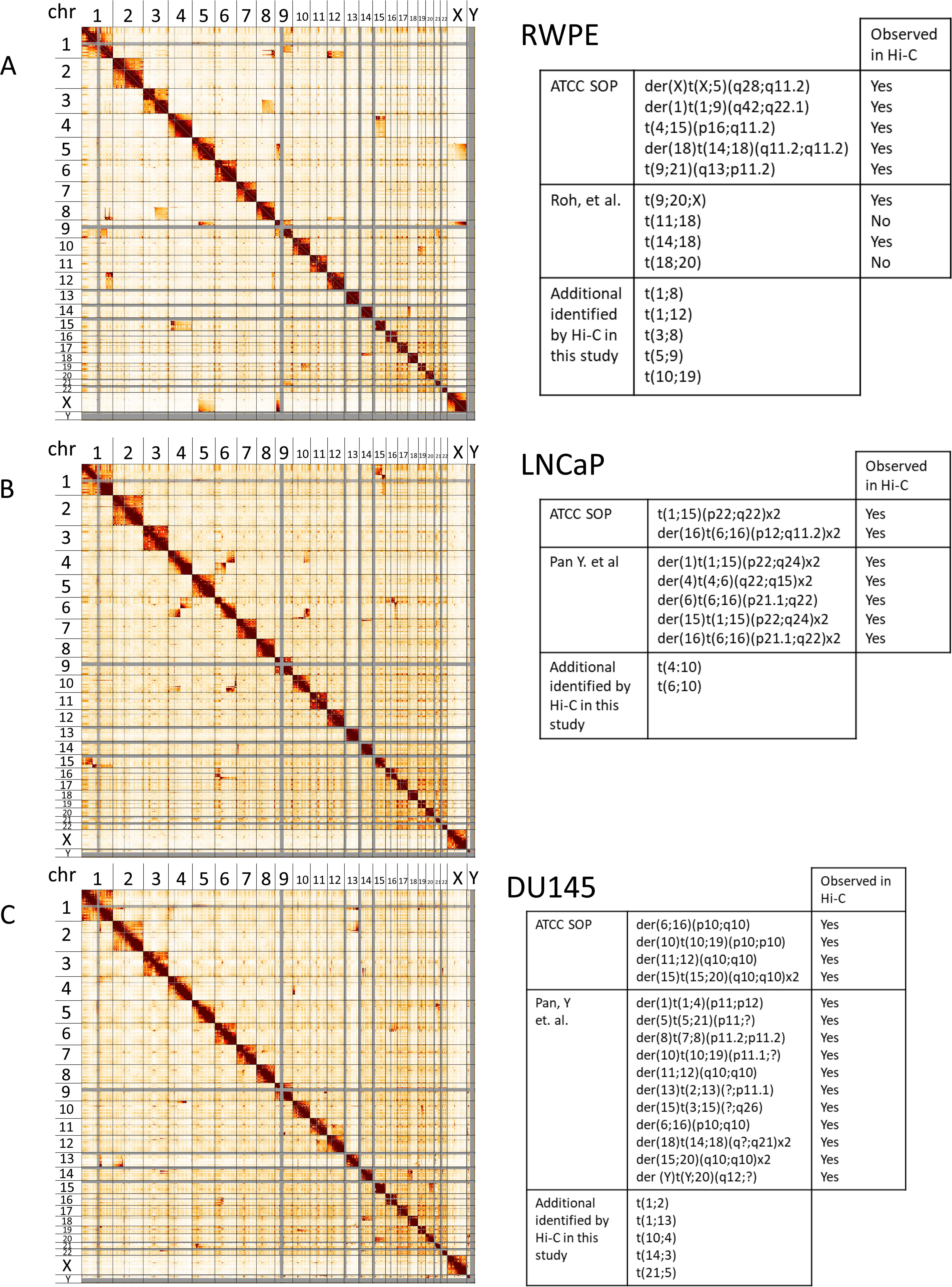
Previously reported translocations for cell lines in this study, as well as some novel ones, are evident in heatmaps of 2.5 MB resolution Hi-C data. 2.5 Mb Hi-C heatmaps for RWPE (A), LNCaP (B), DU145 (C), 22RV1 (D), PC3 (E), MDAPCa2a (F), MDAPCa2b (G) and VCaP (H). Translocations between chromosomes appear as high interaction frequency areas away from the diagonal. Translocations previously described by SKY analysis are listed on the right of each heatmap, along with those events identified in this study; our study has an 83% concordance with previously reported SKY data. The inherent higher resolution of the data obtained by Hi-C makes it possible to identify smaller translocation events.

**Supplementary figure 3.**
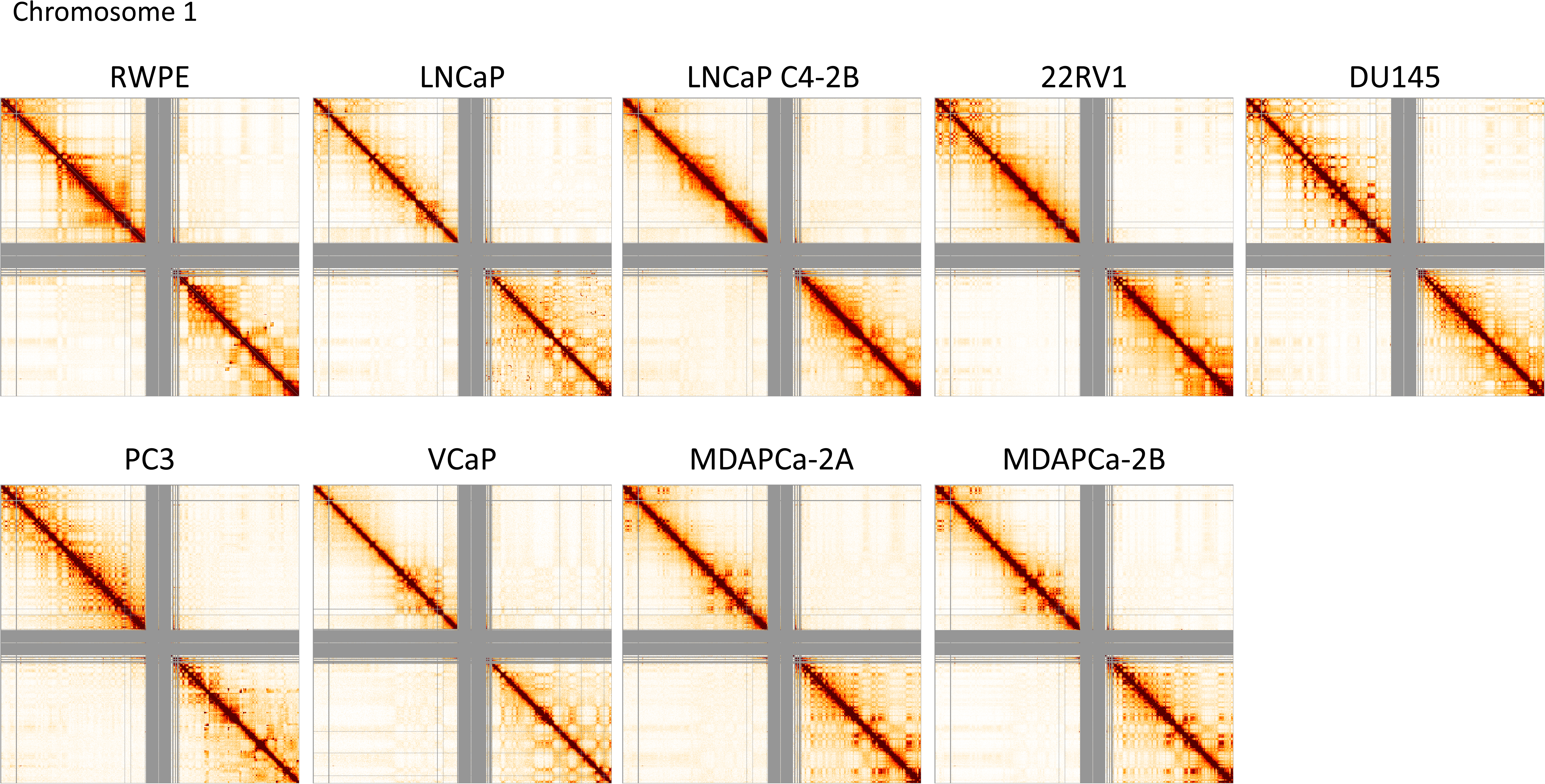
Hi-C Heatmaps for all chromosomes, per cell line, at a 250 kb bin resolution.

**Supplementary Figure 4.**
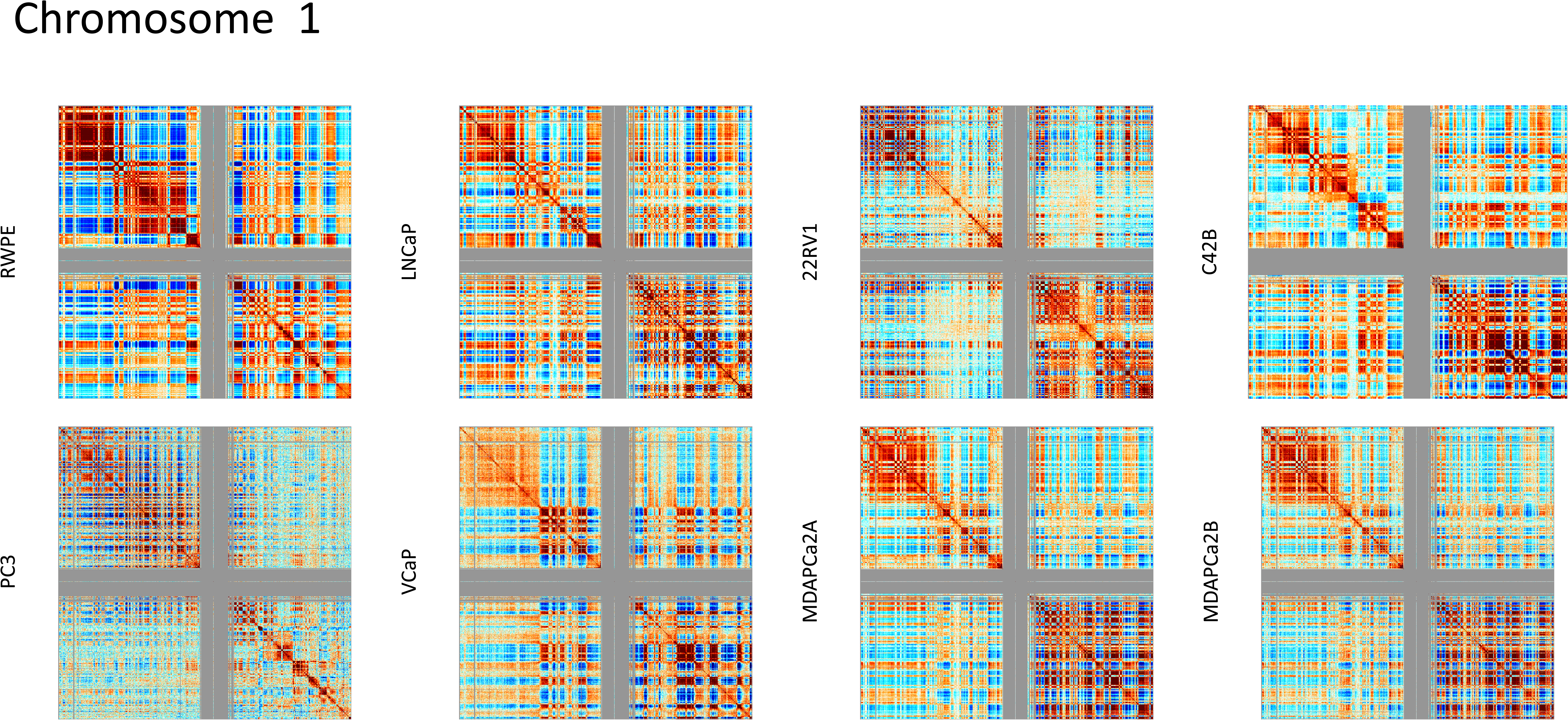
Cis Interaction heatmaps of 250kb-binned data, per chromosome, per cell line.

**Supplementary Figure 5.**
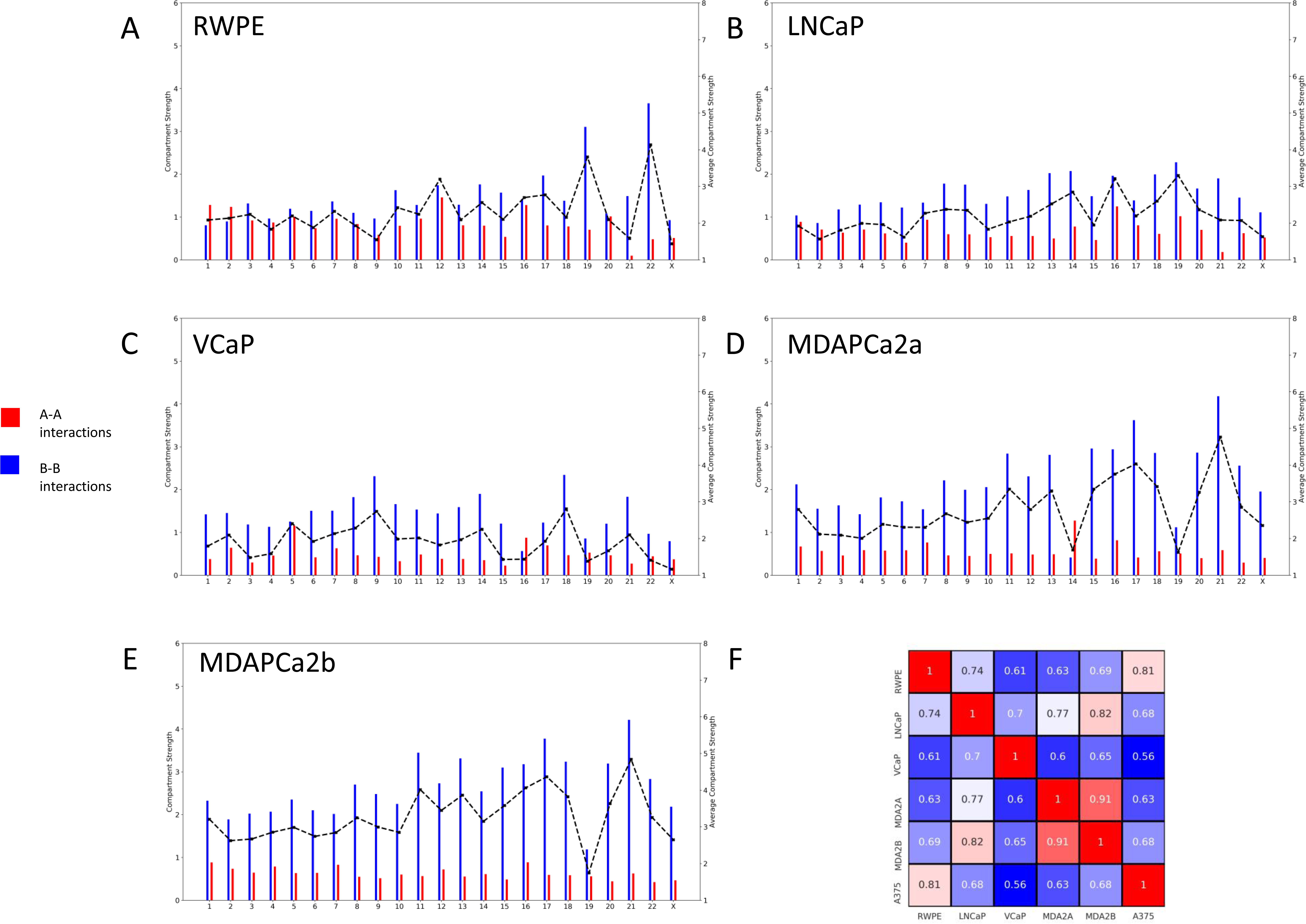
Compartment strength and similarity across cell lines. (A-E) Each graph shows the A-A interaction strength (red) and B-B interaction strength (blue) within each chromosome for the indicated cell line. The dotted line represents the average compartment strength. Compartment strength calculation is defined in Methods. (F) Pearson correlation coefficient between the PC1 compartment tracks genome-wide for the indicated cell lines. A375 is included as a different epithelial-derived cancer (melanoma).

**Supplementary Figure 6.**
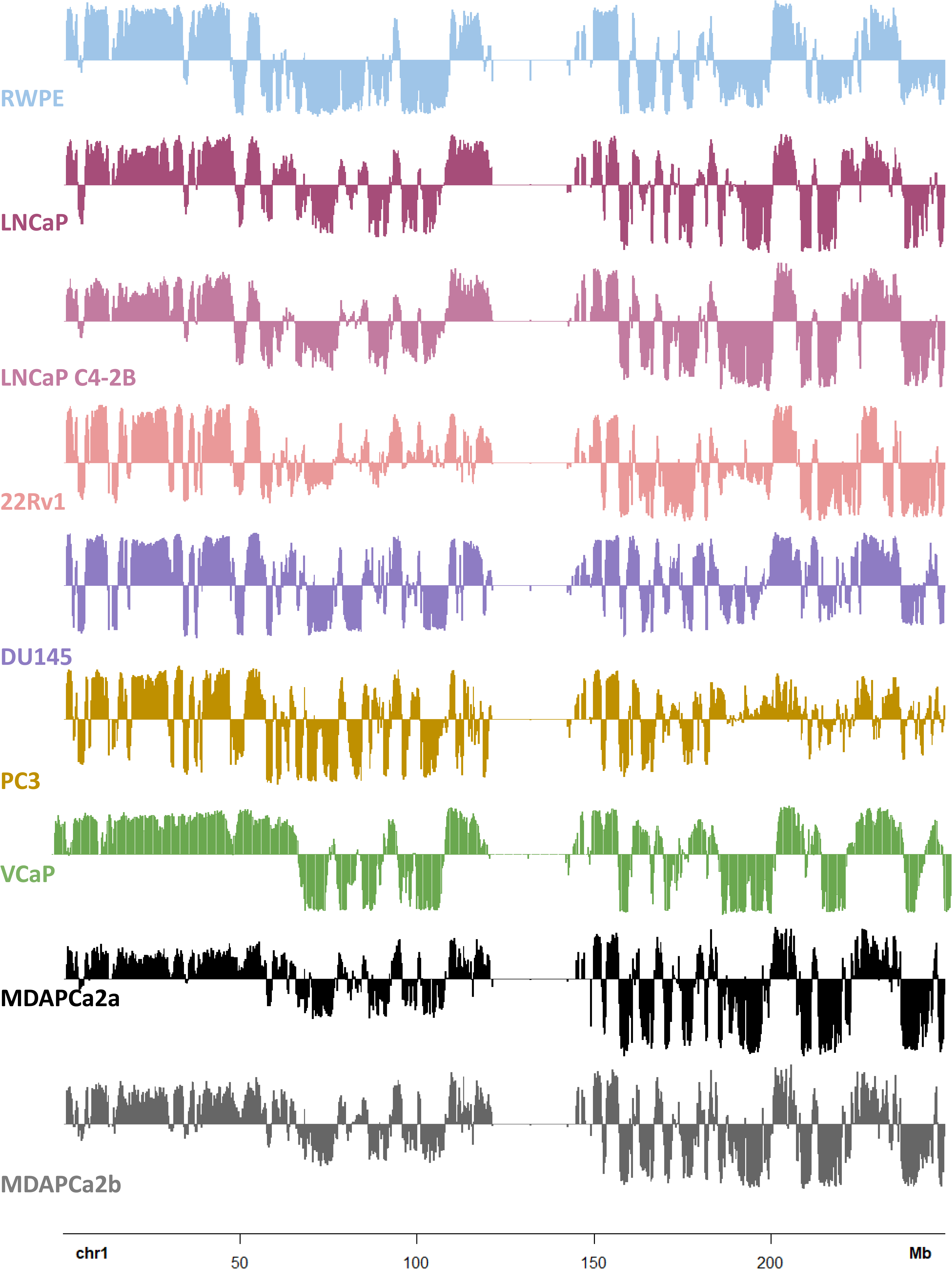
Plots of the 1st eigenvector for each chromosome, obtained from principal component analysis (PC1) of 250 kb-binned data, reveal the compartment identity. Compartment tracks, per chromosome for all the cell lines in this study. From the top: RWPE (light blue), LNCaP (violet), LNCaP C4-2B (pink), 22RV1 (coral), DU145(purple), PC3 (gold), VCaP (green), MDAPCa2a (black) and MDAPCa2b (grey). The high concordance of compartmentalization among tracks for all cells in our model for progression suggests that compartment switches are a consequence of intended biological function.

**Supplementary Figure 7.**
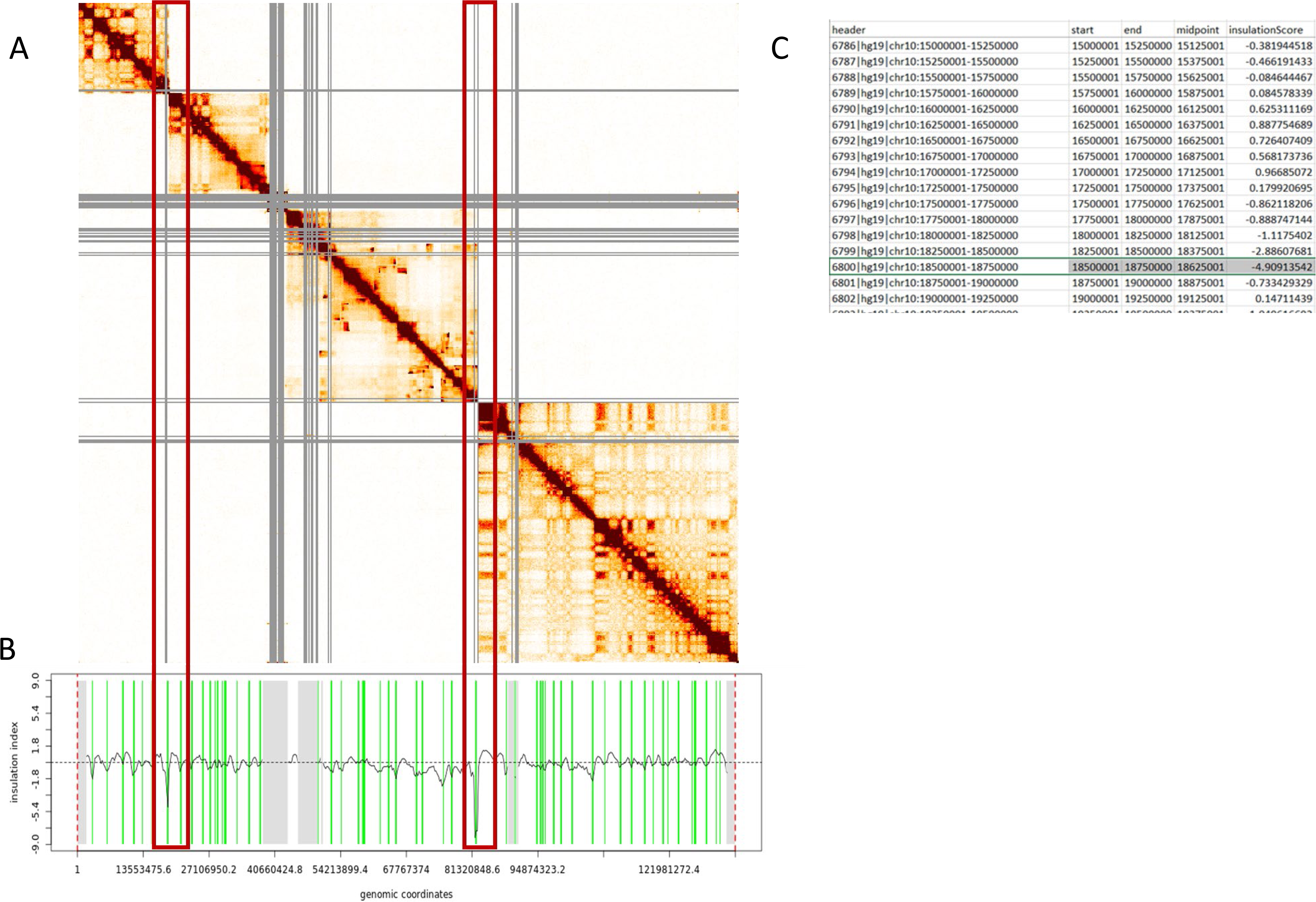
Mathematical reconstruction of highly segmented genomic regions for compartment analysis. A) Hi-C Heatmap of a highly segmented chromosome (chr10, PC3). This type of fragmentation precludes compartment analysis. B) By performing the insulation calculation on the 250 kb-binned data, using an insulation square size of 1.5 Mb it is possible to identify the largest dips in insulation plot, which denote the breaks in the chromosome. C) Compartment analysis is performed on the fragments defined by the bin location. For that particular chromosome, the compartment analysis for RWPE is done for the fragments, to provide a fair comparison for subtraction.

**Supplement Figure 8.**
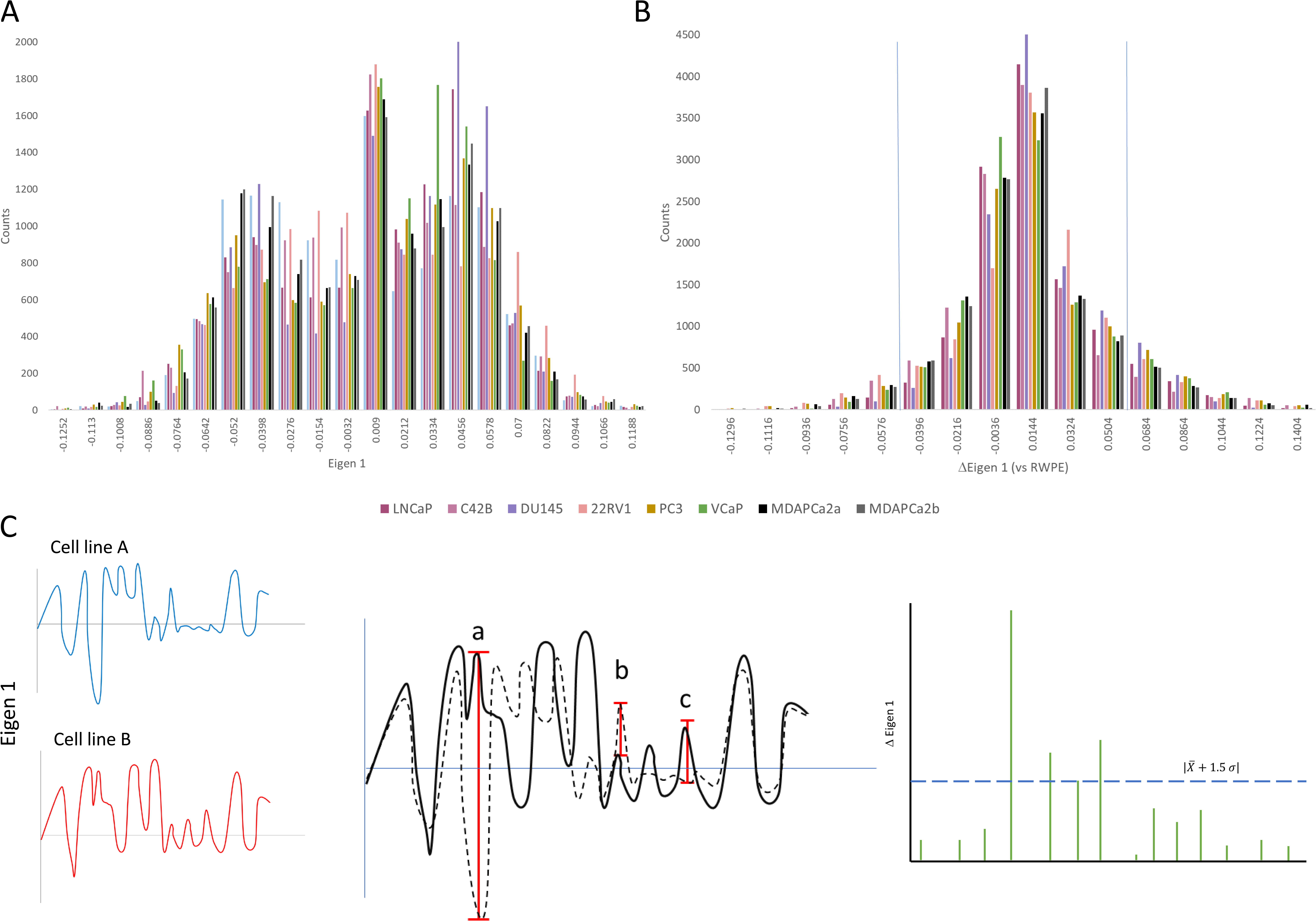
Analytical strategy to determine compartment change relevance in Prostate cancer progression. A) A histogram of the genome wide PC1 values from compartment analysis, per bin at 250kb resolution shows a trimodal distribution for all cell lines in our model for progression. High, positive values represent the A compartment and low negative values represent the B compartment. The third mode represents enrichment of values close to zero. B) A per-bin subtraction of the PC1 values from RWPE creates a distribution closer to normal. Changes in PC1 that exceeded +/- 1.5 of the standard deviation were considered significant compartment changes for the purposes of this study. C) A graphical representation of the ΔPC1 analysis for identification of compartment switches. When superimposing two Eigen 1 tracks, we observe three possibilities: a) a clear, significant compartment change, b) a slight change in the same compartment or c) a modest change in compartments. Normalization against RWPE and thresholding is a straightforward way to identify candidate genomic bins for compartment switches.

**Supplementary Figure 9.**
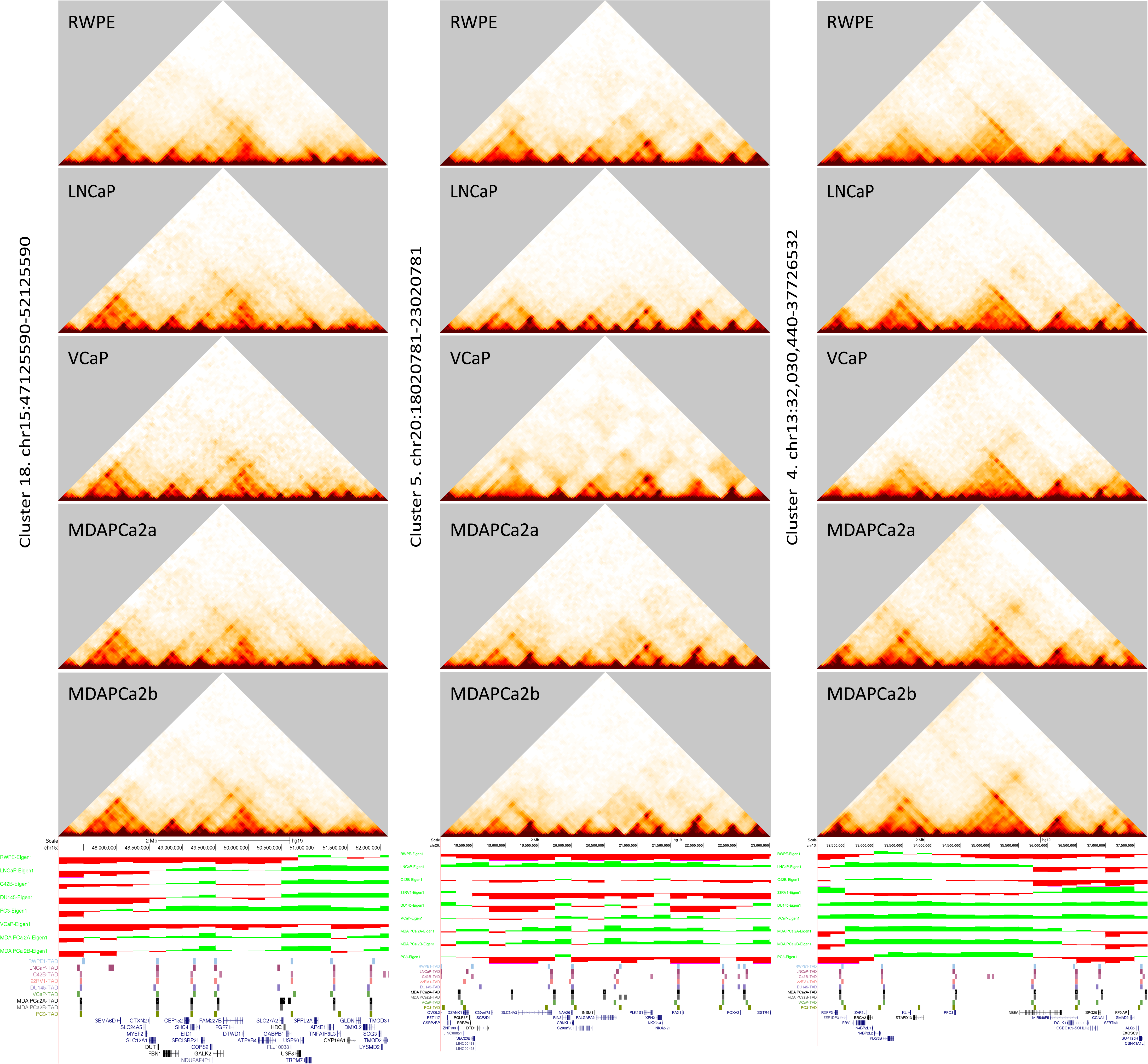
Examples of gene clusters Examples of genes that switch compartments. Left. Cluster 18 includes the following genes: FBN1, CEP152, SHC4, EID1, SECISBP2L, COPS2, NDUFAF4P1, GALK2, FAM227B, FGF7, DTWD1, ATP8B4, SLC27A2 and HDC. While in some cell lines the whole area switches compartments, in others the B compartment persists. Center. Cluster 5 includes the following genes: Cluster with SLC24A3, RIN2, NAA20, CRNKL1, CFAP61, INSM1, RALGAPA2, PLK1S1 XRN2, NKX2-4, NKX2-2, PAX1, FOXA2 and TTC6. Changes to the A compartment in all cell lines in the main progression axis. Right. Cluster 4 changes compartments as a cluster (in DU145, VCaP and MDA lines) or in fragments. It comprises the following genes: Fry (switches B to A) BRCA2 (switches to the B compartment in 22RV1), and N4BP2L1, RFC3, DCLK1, CCNA, SMAD9, ALG5, ZAR1L, RXFP2, and EEF1DP3, all of which switch from the B to the A compartment

**Supplementary Figure 10.**
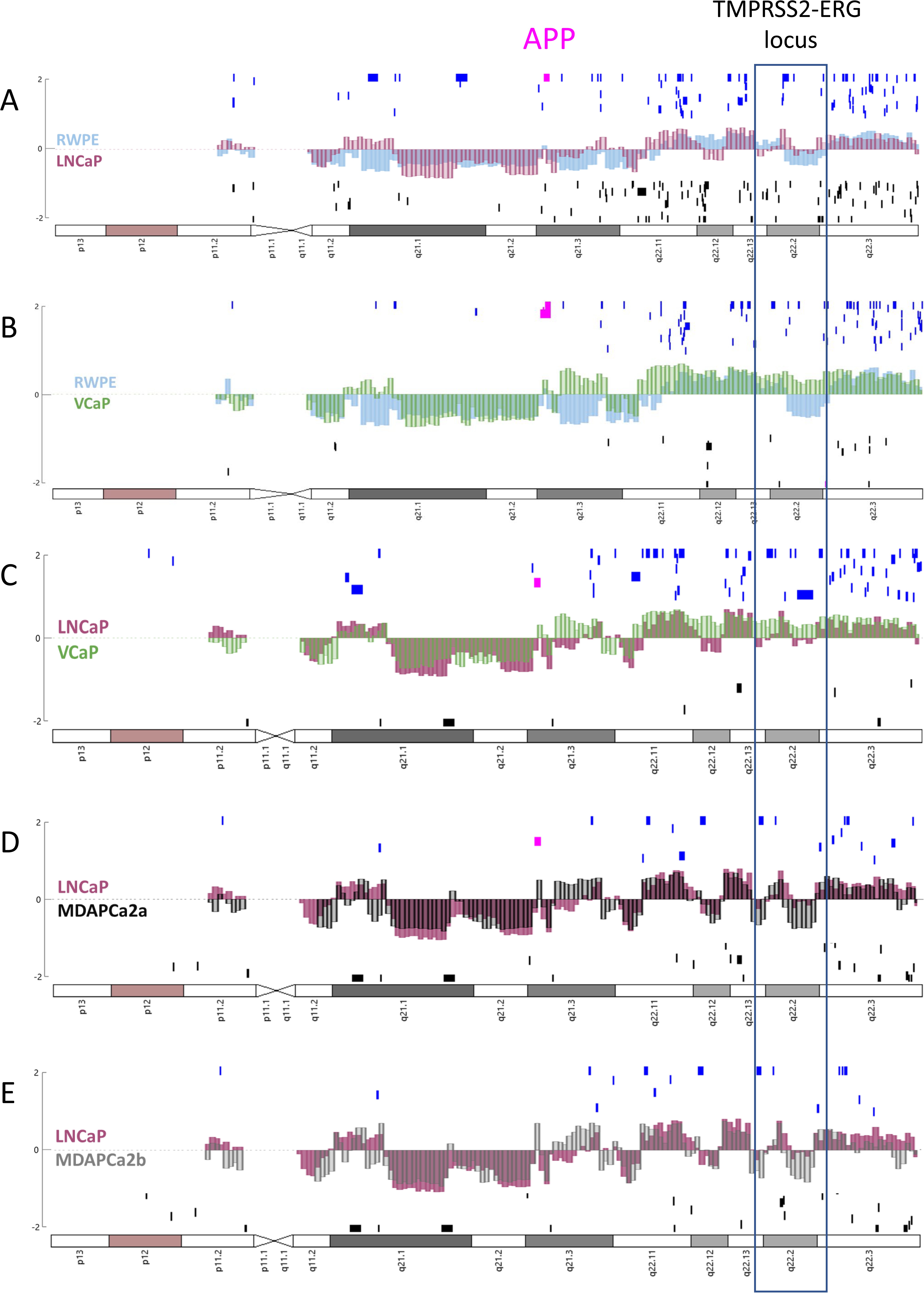
The amyloid precursor protein (APP) – beta secretase 2 (BACE2) axis, located in chromosome 21 is a potential connection between transcriptional activity and compartment switching. Compiled data aligning microarray expression (Log2) for genes along chromosome 21. Genes whose expression increases ≥2 fold or decreases ≤2 fold in the comparison, with a p value <0.05 represented as vertical tick marks along the X axis (chromosome location). Chromosome compartment tracks overlay, superimposed along the microarray track). APP-specific probes highlighted in pink. An increased APP expression is observed in all comparisons, except for MDAPCa2b/LNCaP. A, B, C) Data from HG-U133_Plus_2 microarray comparing LNCaP/RWPE, VCaP/RWPE and VCaP/LNCaP, respectively D,E) Data from Clariom S microarray, comparing MDAPCa2a/LNCaP and MDAPCa2b/LNCaP, respectively

**Supplementary Table 1.**
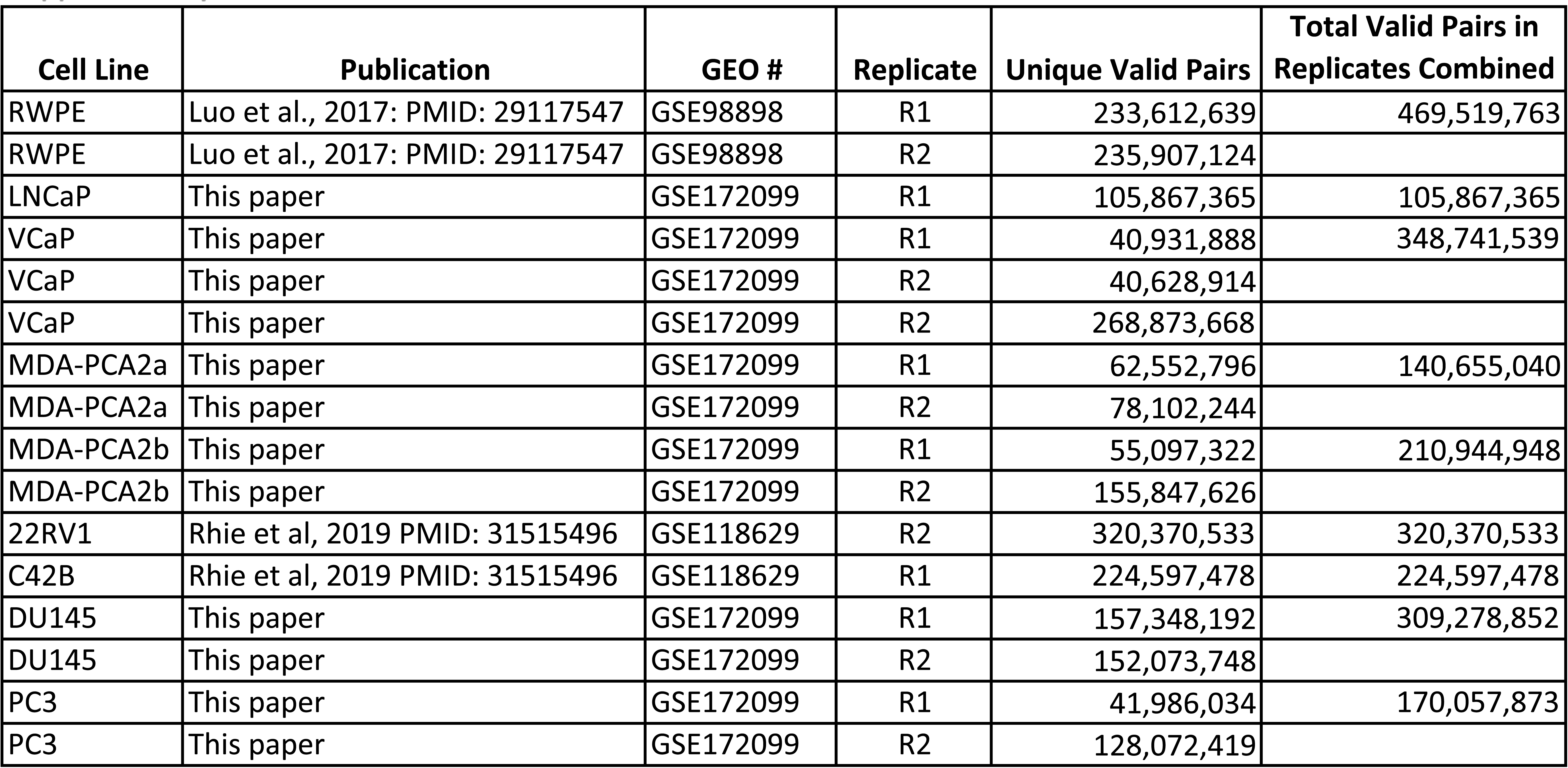
Database references and valid pair counts for all datasets used in this study.

**Supplementary Table 2.**
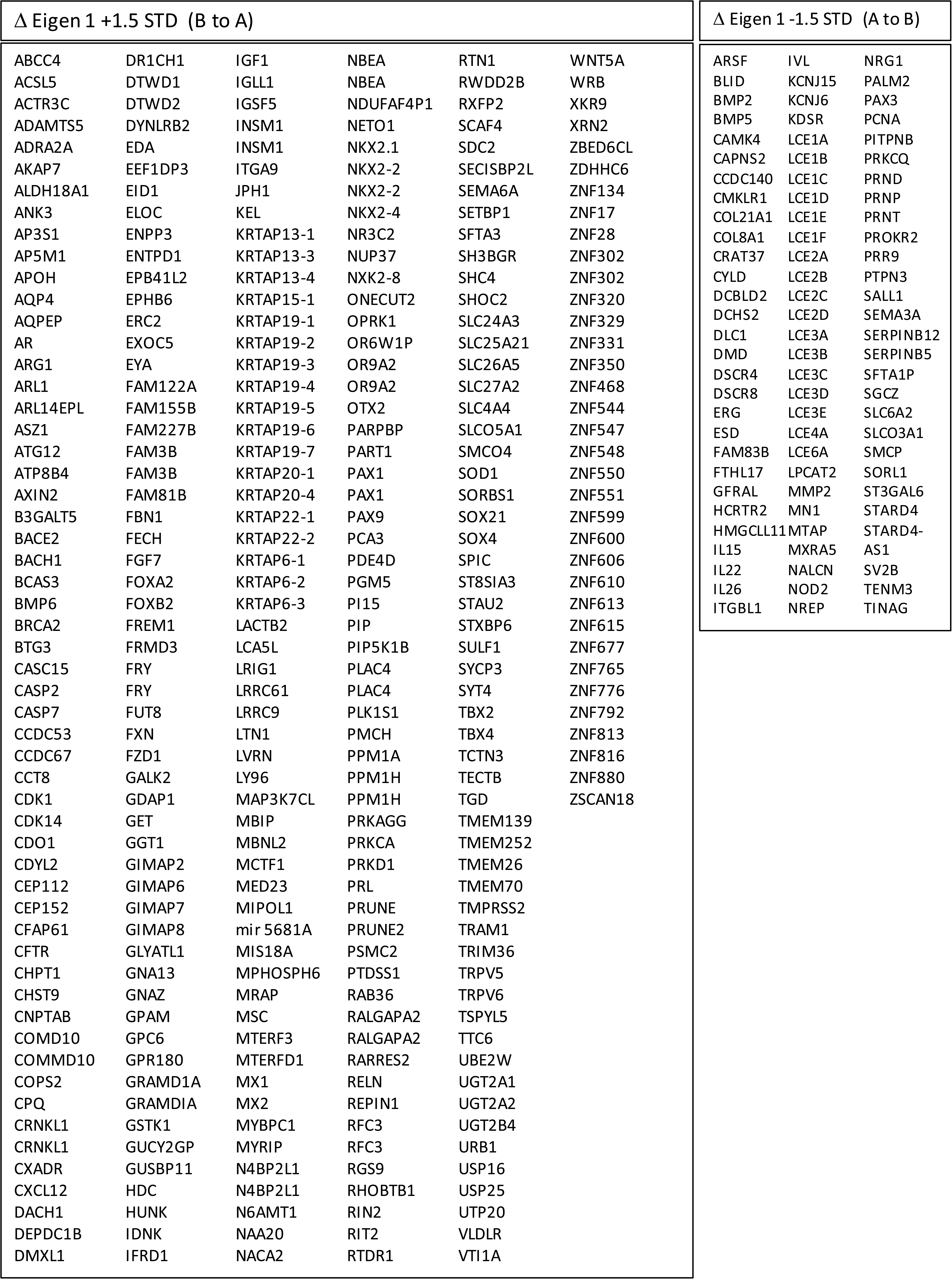
Genes identified through ΔEigen 1 analysis (In at least 6 cancer cell lines OR in all 4 cancer cell lines represented in the axis). Eighty five percent of all genes identified as switching from the B to the A compartment are included in 48 proximal clusters. Of the genes that switch from the A to the B compartment, 74% are included in 16 clusters.

**Supplementary Table 3.**
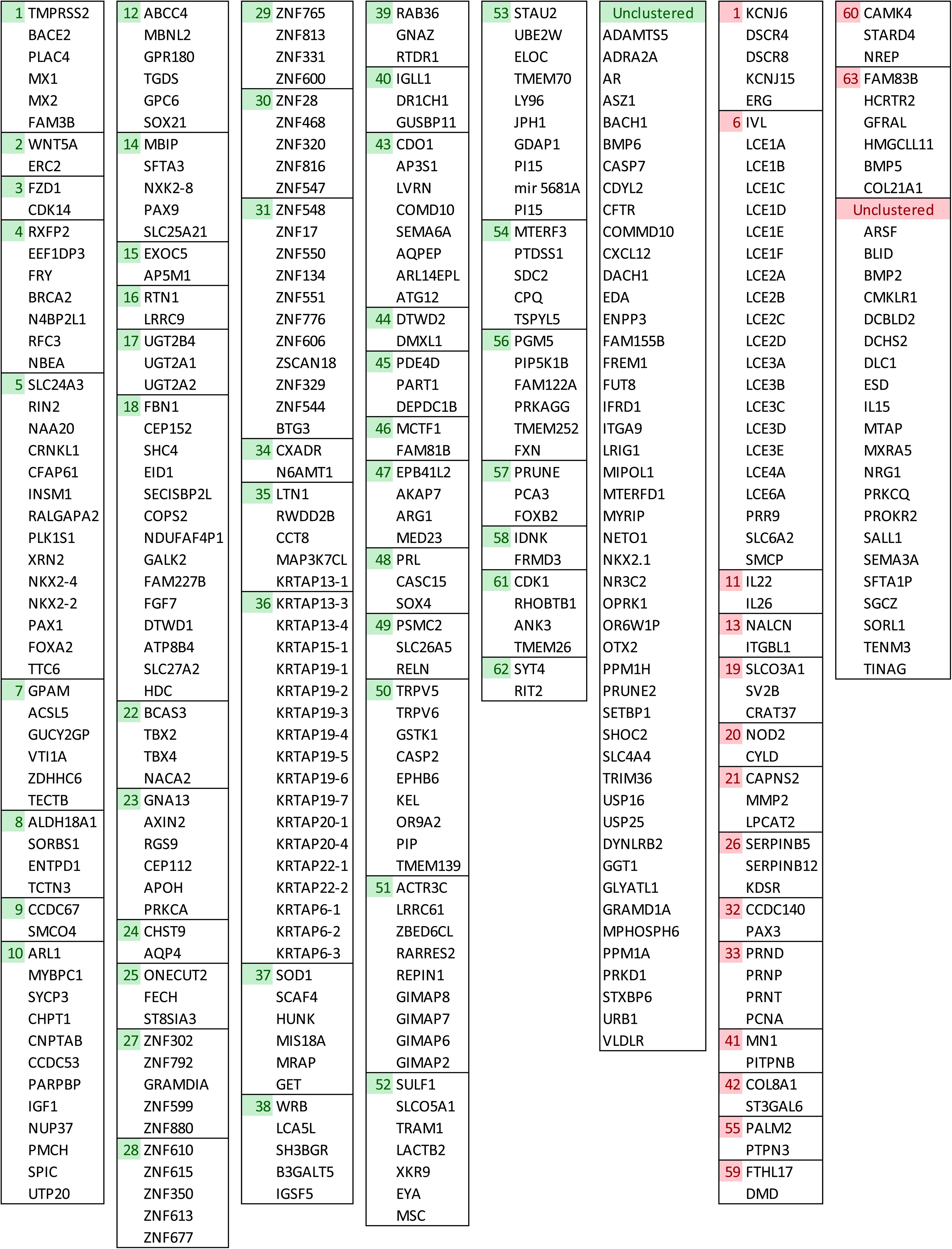
**Gene clusters that switch compartment identity with progression** Gene clusters that switch from the B to the A compartment are highlighted in green, those that switch from the A to the B compartment, in red.

